# Independent roles of Arp2/3 complex and RIC4 protein in the control of epidermal cell shape

**DOI:** 10.1101/2025.09.10.675338

**Authors:** Judith García-González, Lenka Havelková, Erica Bellinvia, Israel Corrales, Přemysl Pejchar, Martin Potocký, Kateřina Schwarzerová

## Abstract

The aim of this study was to determine whether RIC4, a CRIB domain-containing protein that functions as an effector of ROP GTPases, and the Arp2/3 complex, an actin nucleator, functionally cooperate in controlling the shape of Arabidopsis cotyledon epidermal cells. The combination of KO mutants demonstrated that loss of RIC4 and loss of the Arp2/3 complex result in completely opposite epidermal cell shape phenotypes. The double KO mutation is phenotypically similar to the Arp2/3 mutation, and the effect of RIC4 loss is completely eliminated. Analysis of overexpression revealed that excess RIC4 significantly suppresses the formation of pavement cell lobes. However, RIC4 does not require an active Arp2/3 complex for this effect. Our data further show that overexpression of RIC4 has a specific actin stabilization effect in cotyledon epidermal cells. However, we could not demonstrate that actin stabilization is directly related to the cell shape changes that RIC4 overexpression induces. In conclusion, RIC4 and the Arp2/3 complex do not share the same signaling pathway in their function in the control of cotyledon epidermal cell shape.

---

Epidermal cells of the leaves of many plants adopt a distinctive shape resembling a jigsaw puzzle, which is thought to enhance mechanical stress resistance across the tissue (Sapala et al. 2018). This epidermal cell shape is the result of a morphogenetic process that is controlled at several levels. Current research suggests that an interconnected complex of control mechanisms is involved in the initiation and growth of these puzzle-like cells, including signaling at the plasma membrane, formation of membrane microdomains, reorganization of the cytoskeleton, and subsequent synthesis of a cell wall with domain-specific mechanical properties (Liu et al. 2021). While many of the components of the control mechanism are well-defined in terms of their contribution to epidermal cell morphogenesis, they have yet to be integrated into a cohesive, experimentally supported model that explains the full patterning of cell-wall domains. The role of microtubules in shaping epidermal cell shape is probably best explored. Cortical microtubules guide the direction of deposition of cellulose microfibrils (Paredez et al. 2006). These microfibrils form the primary load-bearing framework of the cell wall, creating anisotropic mechanical properties, where regions with dense, aligned cellulose resist turgor-induced expansion (Baskin 2001). According to the main current model, the cell wall with non-uniform mechanical properties with patterned stiffness causes non-uniform expansion under turgor pressure: stiffer zones bulge less, while more permissive regions expand outward, giving rise to the characteristic lobed morphology (Baskin 2005; Szymanski 2014). Published observations suggest that specific domains on anticlinal walls are enriched with cortical microtubules (Fu et al. 2005; Armour et al. 2015; Bidhendi et al. 2019; Altartouri et al. 2019; Lauster et al. 2022), although there is still doubt about the exact localization of microtubules in a given cell (Liu et al. 2021) and other studies have not consistently confirmed this arrangement (Zhang et al. 2011; Belteton et al. 2018). The cortical microtubules in the enriched domains would deposit more cellulose at this site and locally stiffen the cell wall, which gives rise to indentations in pavement cells. Pectins—particularly demethylesterified homogalacturonan— also play a critical role by accumulating at the future neck regions of anticlinal walls, to locally increase stiffness prior to visible lobe formation. Furthermore, arrays of pectin nanofilaments in these walls create a spatially patterned, reversible context that restricts neck expansion while permitting lateral stretching at the emerging lobes (Majda et al. 2017; Altartouri et al. 2019; Haas et al. 2020; Lin et al. 2022).

The organization of cortical microtubules is controlled by signaling pathways involving the Rho of Plants (ROP) GTPases (Smokvarska et al. 2021). The formation of alternating lobes and indentations is coordinated by the spatially segregated activity of ROP2/4 and ROP6 (Fu et al. 2005, 2009). It is well established that ROP6 is localized to the plasma membrane at indentations, where it promotes the activity of its effector, the CRIB domain-containing protein RIC1 (Fu et al. 2009). There, RIC1 interacts with the microtubule-associated protein katanin to induce microtubule ordering, supporting indentation formation in these regions (Lin et al. 2013). The ROP6 signaling pathway at indentations actively suppresses ROP2/4 activity in those regions, establishing complementary domains for cell-shaping signals. This inhibition is executed via the microtubule-dependent recruitment of PhGAP proteins to these sites, thereby inactivating ROP2 (Lauster et al. 2022). Moreover, phosphorylation of PhGAPs by the kinase BIN2 stabilizes their localization in indentation domains, ensuring sustained ROP2 inactivation and robust domain separation (Zhang et al. 2022). At the plasma membrane of lobes, ROP2/4 further reinforces the maintenance of alternating domains by inactivating RIC1, preventing microtubule organization and allowing cell wall loosening for correct outgrowth (Fu et al. 2005, 2009).

While the link of the signaling pathway using ROP6 to the organization of microtubules at anticlinal cell walls in indentations is well characterized, events in lobes domains are not as well defined. Previous work has suggested that the lobes-located ROP2/4 signaling pathway, in addition to restricting MT formation through sequestration of RIC1 (Fu et al. 2005), also activates the formation of fine actin in lobe tips (Fu et al. 2002) through interaction with its effector RIC4 (Fu et al. 2005; Xu et al. 2010, 2014). Indeed, it is indisputable that the actin cytoskeleton also plays an important role in cell patterning, as latrunculin B treatment (Akita et al. 2017) and a number of actin mutants (Mathur et al. 2003; Le et al. 2003; Li et al. 2004; Zhang et al. 2005; Kotchoni et al. 2009; Rosero et al. 2016; Cifrová et al. 2020; Bellinvia et al. 2023) exhibit pavement cell shape phenotypes. The model predicts that actin filaments in the lobes promote vesicle transport of cell wall components such as hemicelluloses, pectins or their modifying enzymes to enable cell wall expansion (Fu et al. 2002, 2005; Breuer et al. 2017; Chebli et al. 2021). Actin accumulation in lobes has been suggested in works that primarily used transient mTalin expression or fixed material for actin labelling (Fu et al. 2002; Frank and Smith 2002; Li et al. 2003; Panteris and Galatis 2005; Djakovic et al. 2006; Xu et al. 2010). However, actin accumulation in the lobes of growing epidermal cells was not observed using stably expressed in vivo markers (Zhang et al. 2013; Armour et al. 2015; Lauster et al. 2022), and some work suggests that actin in lobes may occur as a consequence of being displaced by cortical microtubules from sites of indentation. Thus, the specific accumulation of actin in expanding lobes may be an observation that needs to be confirmed. Moreover, an experimentally confirmed link between ROP2 and actin in this model is still lacking, as the frequently mentioned RIC4 protein does not directly interact with actin (Fu et al. 2005; Lin et al. 2015). Finally, the function of polymerized actin itself in controlling lobe expansion also remains unknown.

Nevertheless, it is clear that actin plays a specific role in the morphogenetic process of leaf epidermal cells. One piece of evidence for the influence of actin on this process is provided by some actin mutants in which phenotypes associated with epidermal patterning are well known and documented. Among the most thoroughly researched actin-related mutants impacting the shape of pavement cells are those in the Arp2/3 complex, which is one of the only two known actin nucleators in plants. The Arp2/3 complex needs activation by the SCAR/WAVE complex to bind actin and initiate the formation of additional actin filaments (Weaver et al. 2003; Frank et al. 2004). Mutants lacking a functional Arp2/3 or SCAR/WAVE complex have an impaired ability to form lobes (Mathur et al. 2003; Li et al. 2003, 2004; Le et al. 2003; Brembu et al. 2004; Saedler et al. 2004; Zhang et al. 2005, 2013; Djakovic et al. 2006; Kotchoni et al. 2009; Bellinvia et al. 2023). Importantly, the Arp2/3 complex is a potential link between actin and ROP signaling. Subunits of the SCAR/WAVE complex interact with ROPs (Basu et al. 2004, 2008; Uhrig et al. 2007; Feiguelman et al. 2018). In addition, the SCAR/WAVE complex interacts with SPIKE1, a DOCK-family ROP-GEF, which interacts with several ROP GTPases primarily in the ROP-GDP form (Basu et al. 2008). These findings position the Arp2/3 complex downstream of ROP signaling during epidermal cell morphogenesis.

However, it is unclear whether the function of RIC4 in the reorganization of the actin cytoskeleton in epidermal cell lobes is functionally connected to the Arp2/3 complex (Fu et al. 2005). In this study, we use genetic and biochemical tools to confirm or refute the theory that RIC4 and Arp2/3 operate in the same pathway in Arabidopsis epidermal cell formation.

## Material and methods

### Plant cultivation

Plants were grown in peat pellets or in vitro (vertical agar plates containing half-strength Murashige and Skoog medium supplemented with 1% w/v sucrose) under a photoperiod of 16h light:8h darkness and 23 °C and light intensity 110 µmol/m2/s. Arabidopsis thaliana genotypes used in this study were Col-0 (wild-type, wt), *arp2* (SALK_077920.56.00), *arpc5* (SALK_123936.41.55), *arpc4* (SALK_013909; Pratap Sahi et al. 2017), *ric4* (SALK_015799), *phgap1 phgap2* (Stöckle et al. 2016) and *CA-rop2* (Li et al. 2001). *ric4* line was genotyped to confirm insertion using gene and SALK specific primers (**Table 1**). PCR product sequencing confirmed T-DNA insertion after the nucleotide 164 of the coding sequence, corresponding to AA 60 in protein sequence resulting in a stop codon in position AA 80. qRT-PCR was performed to confirm the knock-out state of the line (primers described in **Table 1, supplementary figure 1A**). Briefly, two non-overlapping primer pairs were designed within the RIC4 coding sequence on either side of the T-DNA insertion mapped after nucleotide 164. The pair RIC4-A amplifies a 106bp region upstream of the insertion, and pair RIC4-B amplifies a 124bp region downstream of the insertion. The upstream amplicon (RIC4-A) controls for residual transcript, whereas the downstream amplicon (RIC4-B) tests for read-through across the disrupted locus. The RIC4-A reverse primer and the RIC4-B forward primer span exon–exon junctions, preventing amplification from intron-containing genomic DNA, ensuring that cDNA specificity. Data was analyzed using the protocol described by Ganger and colleagues (Ganger et al. 2017).

**Table 1:**
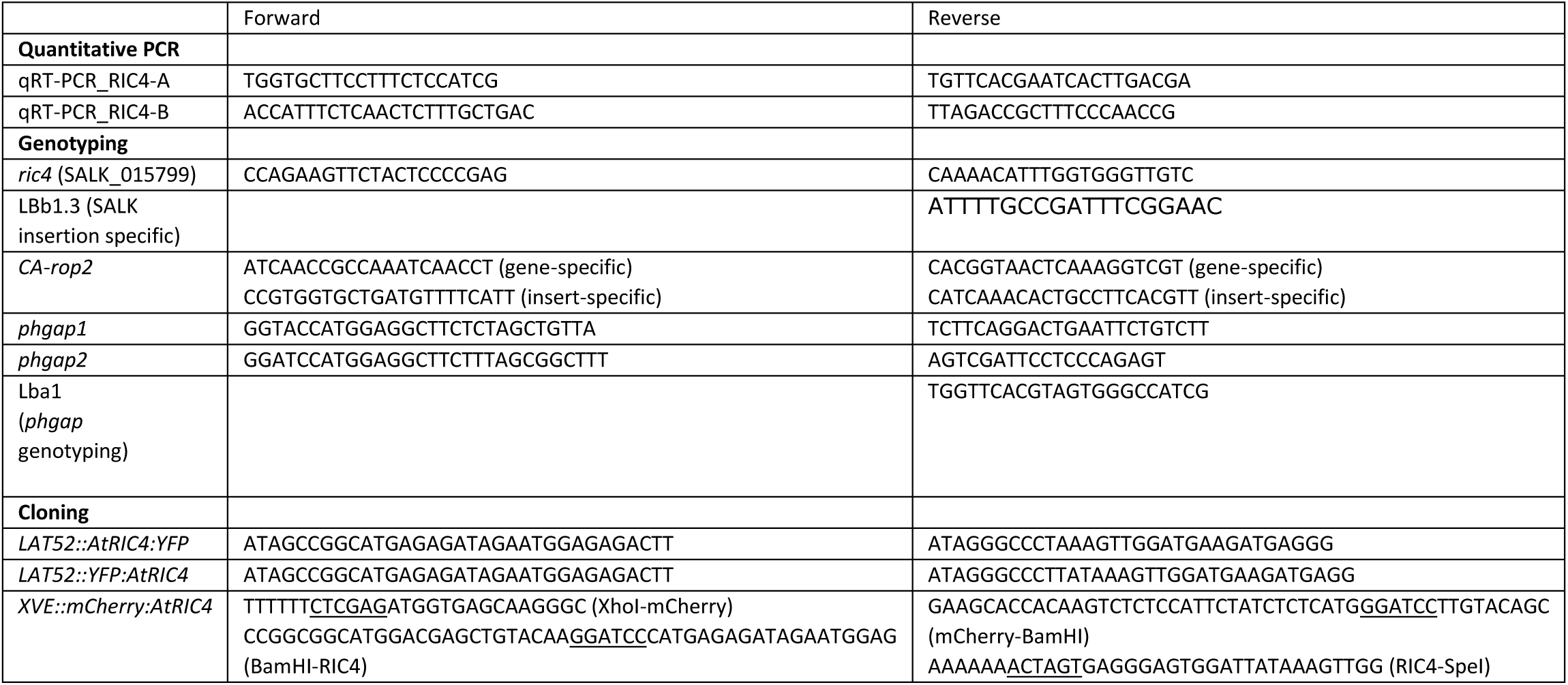
Oligonucleotides used in this study. All sequences are written in 5’ → 3’ sense. For quantitative PCR, two independent *RIC4* primer pairs were used on cDNA, upstream (RIC4-A) and downstream (RIC4-B) of the T-DNA insertion site. For genotyping of homozygous *ric4* (SALK_015799) line, gene-specific primers were used either together to detect the wild-type allele or in combination with the SALK left-border primer (LBb1.3) to detect the insertion allele. The *CA-rop2* line contains a mutated coding sequence in the wild-type genome; accordingly, the gene-specific primer pair yields a single product from the endogenous locus in both wild-type and *CA-rop2* line, whereas the insert-specific primer pair yields two products for the mutant – a shorter band from the intronless cDNA and a longer band from the intron-containing gDNA, allowing for the detection of the presence of the insert. Primer pairs specific to *phgap1* and *phgap2* lines were used in combination with the left border primer Lba1 for T-DNA insertion line homozygote genotyping. Primers for cloning were used as described in the methods; underlined nucleotides denote restriction sites used later for ligation.

### Cloning

To prepare the *Lat52::AtRIC4:YFP* and *Lat52::YFP:AtRIC4* constructs, the RIC4 coding sequence was amplified from *Arabidopsis thaliana* Col-0 cDNA using Q5 High-Fidelity DNA Polymerase (NEB), with primers flanked by NgoMIV and ApaI restriction sites (**Table 1**). The amplified fragments were cloned into the multiple cloning sites of the pollen-specific expression vectors pHD32 (C-terminal) and pWEN240 (N-terminal) (Klahre et al. 2006).

To generate the *XVE::mCherry:RIC4* reporter, total RNA was isolated from wild-type Arabidopsis using the NucleoSpin RNA Plant kit (Macherey-Nagel), and first-strand cDNA was synthesized using M-MuLV reverse transcriptase (Thermo Fisher). The coding sequences of mCherry and RIC4 were PCR-amplified using Q5 High-Fidelity DNA Polymerase (NEB), with primers containing XhoI/BamHI sites for *mCherry* and BamHI/SpeI sites for *RIC4* (see **Table 1**). Amplified products were purified via agarose gel extraction and ligated sequentially into the pER8 vector backbone harboring the XVE estrogen receptor-based transactivator (Zuo et al. 2000). Proper insertion orientation and the integrity of the resulting *XVE::mCherry:RIC4* were confirmed by sequencing.

### Transformation of Arabidopsis

Four to five-week-old plants were transformed according to the modified floral dip method (Zhang et al. 2006). T2 progeny of independent transformants were tested for the expression under induction conditions (1 μM β-estradiol; Sigma-Aldrich, Cat. No. E2758, stock solution 20 μM in DMSO) and representative lines were used in further experiments.

### Transient pollen expression

*Nicotiana tabacum* pollen transformation and the microscopic analysis of expression of RIC4 fused to YFP and pollen tube growth was performed as described in (Pejchar et al. 2020). Particles for biolistics were coated with 1.5 ug of DNA. For detailed protocol of pollen transformation see (Noack et al. 2019).

### Microscopy

For pavement cell shape analysis, propidium iodide (PI; aqueous solution 0.01 mg/ml) stained cotyledons were observed under the laser scanning microscope Leica TCS SP2 using HC PL APO 20.0x/0.70 IMM/CORR λ_BL_ objective (ex: 488 + 496 nm; em: 593-668 nm). Dual-color imaging of mCherry–RIC4 and Lifeact–GFP was performed on a Leica TCS SP8 with an HC PL APO CS2 63x/1.20 water-immersion objective with excitation and emission set for GFP (ex: 488 nm, em: 493-556 nm and mCherry (ex: 561 nm, em: 588-645 nm). Actin dynamics in Lifeact-GFP-expressing cells was recorded on a spinning disc Nikon Eclipse Ti-E (Yokogawa CSU-X1) microscope, with a Plan Apo VC 60x/1.20 WI DIC N2 objective and Andor Zyla sCMOS camera (ex: 488 nm, ex: 535 nm). Actin analysis was performed on videos with 149 frames and 400 ms frame rate. Cotyledons or hypocotyls from 4-day-old plants expressing Lifeact-GFP, or co-expressing Lifeact-GFP and mCherry-RIC4 were cut, mounted into an observation chamber beneath a thin block of agar cultivation medium for better contact with the coverslip. If required, cells overexpressing mCherry-RIC4 were located in the microscope prior to actin cytoskeleton recording.

Pollen tubes expressing RIC4 fusions with YFP were likewise imaged on the same spinning disk system with a 60x Plan Apochromat 1.20 NA water-immersion objective and Andor Zyla sCMOS camera. YFP was excited at 514 nm and collected through a 542/27 nm bandpass filter (Semrock Brightline). Laser and camera settings were kept constant throughout each imaging experiments, allowing for comparative imaging.

### Method of pavement cell analysis and statistics

For pavement cell shape comparison between mutants, cotyledons of seedlings grown for 14 days were incubated for 20-30 min in aqueous solution of PI. For the analysis the effect of inducible mCherry-RIC4 expression, pavement cells of cotyledons of seedlings grown for 8 days on medium with 1-5 μM β-estradiol were analyzed under the confocal microscope. Only cells expressing mCherry were chosen for further analysis. Mock-treated controls (0.1% DMSO) of the same genotype were stained and imaged in parallel, using exactly the same settings. Pavement cell shape parameters were quantified using ImageJ/Fiji (2.16.0/1.54p) platform (Schindelin et al. 2012). Pavement cell shape analysis was carried out as described in (Pratap Sahi et al. 2017). Statistical analysis of pavement-cell shape parameters (area, circularity and solidity) was performed in R (v4.5.0). Biological repetition was included as a random effect and genotype and/or treatment as fixed effects. To optimize model fit, area was log-transformed, and alternative models (log-LMM, Gaussian GLMM, Gamma GLMM) were compared for goodness-of-fit. For each dataset, the final model was chosen based on AIC, BIC, likelihood-ratio test and model assumptions diagnostics. Because circularity and solidity are bounded between the 0-1 interval, they were analyzed via beta-regression GLMMs, after comparison with other LMM/GLMMs models via AIC and likelihood-ratio tests. Final models were further validated for specific assumptions via appropriate diagnostics (residual, dispersion, collinearity, normality and homoscedasticity checks using the performance or DHARMa (Residual Diagnostics for Hierarchical (Multi-Level/Mixed) Regression Models) package simulations alongside Shapiro–Wilks or Breusch– Pagan tests). Tukey-adjusted pairwise contrasts were extracted using the emmeans package. Data visualization was generated with ggplot2.

All tests were two-sided. When only comparisons between induced and uninduced lines were made, exact p-values are reported. When multiple genotypes were compared within a panel, significance is summarized with letters (CLDs), where groups sharing a letter do not differ at the α specified after multiple-comparison adjustment. We used α < 0.01 for significance in all analyses except cases where higher within-group variance existed despite consistent effect. The significance threshold used is disclosed in every caption.

### Actin dynamics measurement

Actin dynamics was recorded in cotyledon and hypocotyl epidermal cells of four-day-old seedlings. Seedlings were grown on standard medium or, for lines carrying the *XVE::mCherry:RIC4* construct, on medium supplemented with either 0.01% DMSO (mock) or 2 nM β-estradiol. In the β-estradiol–treated samples, only cells exhibiting detectable mCherry fluorescence were selected for further analysis.

Actin dynamics analysis was performed based on the method described by Vidali, L. *et al*. (Vidali et al. 2010). Portions of the final code were adapted from the materials used by Gavrin, A. *et al*. (Gavrin et al. 2020). Raw time-series images were processed in ImageJ/Fiji (2.16.0/1.54p) using a custom script. Background subtraction was carried out using the rolling-ball algorithm (radius = 50 pixels), followed by linear contrast enhancement across the full dynamic range. Photobleaching was corrected via histogram matching. To reduce selection bias, regions of interest (ROI) were defined using a randomly generated grid overlay (with line spacing of 20 µm spacing). ROIs were then manually selected within the grid image; these ROIs were saved as a ZIP file. Due to curvature of the epidermis and focal plane variation, only signal from cells touching the coverslip was retained. Areas lacking clear signal were omitted manually. Pairwise Pearson correlation coefficients (r) between image frames within each ROI were computed using a custom MATLAB based on the corr2 function, adapted from Vidali and colleagues (Vidali et al. 2010).

Data analysis and visualization were performed in R (v4.5.0) using librarian to load stringr (1.5.1), openxlsx (4.2.8), dplyr (1.1.4), tidyr (1.3.1), tidyverse (2.0.0), purr (1.0.4), tiff (0.0.12), ggplot2 (3.5.2), glmmTMB (1.1.11), forecast (8.24.0), and ggforce (0.5.0). File metadata were parsed using custom regular expressions, and low-quality data were filtered prior to analysis. Statistical modelling was carried out following the approach proposed by (Spyroglou et al. 2021), and is briefly described in **Supplementary data 1**.

### Image segmentation and actin structure analysis

The algorithm for actin enhancement, segmentation and organization measurement was implemented in python, based on the work of (Liu et al. 2018, 2019) and (Li et al. 2023). Image enhancement and segmentation method is described in **Supplementary data 2**.

Occupancy was computed as the ratio of actin-positive pixels in the binary mask to the total image area, providing a measure of spatial coverage. Anisotropy was quantified by performing local orientation analysis on the skeletonized structures. Specifically, the structure tensor was calculated within sampling regions across each frame, and anisotropy was defined as the absolute difference between the principal eigenvalues of the tensor. These local anisotropy values were weighted by filament length and averaged across the image to obtain a global alignment score. Skewness was computed from the distribution of intensity values within the actin mask and reflects the asymmetry of filament brightness. The coefficient of variation (CV) was calculated as the standard deviation divided by the mean of the pixel intensities within the same masked regions, providing a measure of local intensity heterogeneity. Frame-wise results were averaged over time for statistical evaluation.

Statistical analyses were conducted in R (v4.5.0). Per-ROI trait means were transformed to meet modelling assumptions—occupancy and anisotropy were scaled to values in the 0-1 interval, while skewness and CV were log-transformed. For each trait we fitted and compared ordinary linear models, linear mixed-effects models, and (where appropriate) generalized linear mixed models by AIC and residual diagnostics; models that failed to improve fit or violated assumptions were discarded.

Occupancy and anisotropy were best described by beta-family GLMMs (glmmTMB) with a logit link, incorporating fixed effects for treatment, genotype and their interaction, plus random intercepts for individual plants and biological replicate. Skewness was analyzed using a robust linear regression on the log-transformed values with the same fixed-effect structure, and CV was modeled by linear regression on log-transformed CV with fixed effects for treatment, genotype and interaction. Pairwise contrasts between DMSO and β-estradiol within each genotype were estimated for the linear and mixed models, and by bootstrap resampling of residuals for the robust skewness model. Final model assumptions were verified via Shapiro–Wilk and Breusch–Pagan tests for the linear models and by DHARMa simulations for the GLMMs.

### *arpc5* and *ric4* mutants have opposite phenotype in pavement cell shape

As the first step of the analysis of the interaction of pathways using the Arp2/3 complex and the RIC4 protein, we carefully compared morphological defects in pavement cells in single and double mutants of Arabidopsis. The shape of adaxial epidermal cells in the *arpc5* line, expressed as circularity and solidity variables as indicator or lobe number and outgrowth, and the size of epidermal cells, expressed as area, confirmed previously published results (Pratap Sahi et al. 2017). In comparison to wild-type (**Fig. 1a**), the loss of Arp2/3 complex (**Fig. 1b**) results in an increase in both circularity and solidity (**Fig. 1 e, g**) as an expression of reduced complexity of epidermal cell shape. Cell surface area was slightly but statistically significantly increased in the *arpc5* mutant (**Fig. 1f**). The cell shape of the *ric4* single mutant (**Fig. 1c**) showed the opposite trend. Loss of RIC4 led to a statistically significant decrease in circularity and solidity compared to control (**Fig. 1e, g**). The cell area of *ric4* was statistically significantly increased compared to wild-type, and the values were comparable to those of *arpc5* (**Fig. 1f**). The *arpc5/ric4* double mutant (**Fig. 1d**) had circularity and solidity values clearly similar to the *arpc5* single mutant with some additive effect (**Fig. 1e, g**) while cell area was comparable to wild-type controls (**Fig. 1f**). The results suggest that the function of the Arp2/3 complex, which is executed during epidermal cell shaping is dominant to that of the RIC4 protein. However, additive changes in both shape and size features indicate measurable interaction elements characteristic of epistasis in complex traits, which suggests that the Arp2/3 complex and RIC4 might operate in parallel but partially overlapping pathways.

**Figure 1:**
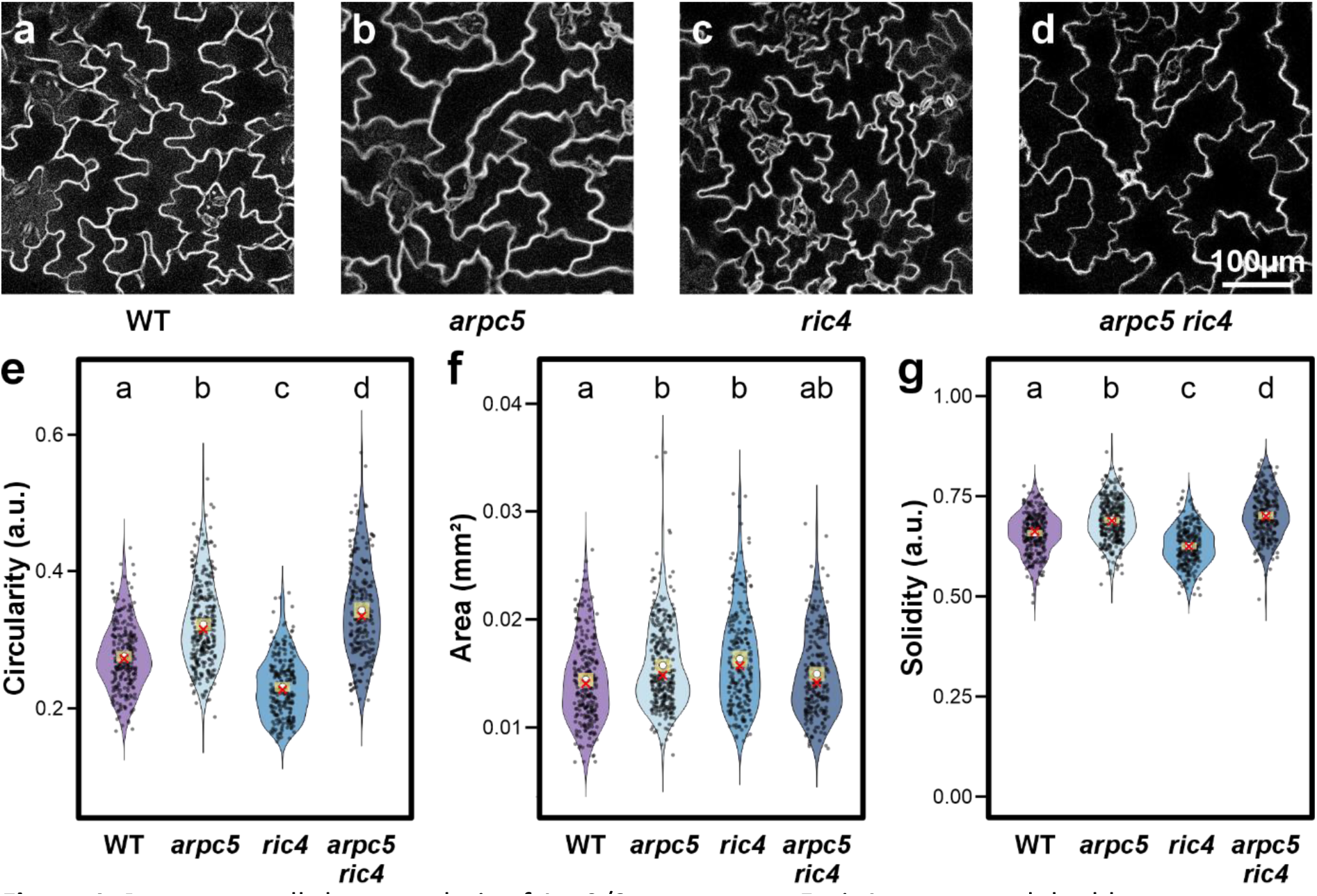
Pavement cell shape analysis of Arp2/3 mutant *arpc5*, *ric4* mutant and double mutant *arpc5/ric4*. (a-d) Representative pictures (maximum-intensity z-projections) of propidium iodide-stained cotyledon pavement cells: wild-type (a), *arpc5* (b), *ric4* (c) and the double mutants *arpc5/ric4*. (e-g) Violin plots of cell circularity (e), area (f) and solidity (g) for each genotype (n=239-314 cells per group). Statistical significance was assessed by mixed-effects (log-transformed area) or beta-regression (circularity and solidity), followed by Benjamini–Hochberg (FDR)-adjusted pairwise comparisons. A two-sided threshold of α = 0.05 (95% confidence level) was used to define significance. Distributions are overlaid with 99% confidence intervals (yellow bar), mean (white circle) and median (red cross). Scale bar = 100 µm.

### Overexpression of RIC4 results in reduction of cell lobes formation

Proteins of the ROP GTPase regulatory pathway belong to multigene families with considerable redundancy, so single loss-of-function mutants often show less pronounced phenotypes, whereas their overexpression typically produce stronger effects. Therefore, we tested the effect of RIC4 overexpression on cell shape in different cell types and its relationship to Arp2/3 mutants. Arabidopsis RIC4 was fused to YFP at either the N- or C-terminus and expressed under the pollen-specific Lat52 promoter in tobacco pollen tubes. Both fusion variants (Lat52::AtRIC4-YFP and Lat52::YFP-AtRIC4) accumulated at the growing tip (**Supp. Fig. 1a, c**), and higher expression levels caused apical swelling, a characteristic effect of RIC4 overactivity (**Supp. Fig. 1b, d**) (Gu et al. 2005). These assays confirmed that our RIC4 fusions are functional. (Gu *et al*. 2005). To analyze RIC4 function in epidermal cells, we cloned the protein into a vector with the estradiol-activated XVE system (Zuo et al. 2000) and Arabidopsis *ric4* plants were stably transformed with the resulting vector harboring the mCherry-RIC4 fusion protein. In comparison to uninduced *ric4* mutant (**Fig. 2a**), transgenic *ric4* plants expressing mCherry-RIC4 **(Fig. 2b)** after induction by 1 μM beta-estradiol had significantly increased cell circularity and solidity (**Fig. 2c, e)**, while their cell area was statistically significantly lower than that of mock-treated plants (**Fig. 2d**). Three independent transformation lines showed the same trend (**Supp. Fig. 2**). We concluded that overexpression of RIC4 causes the opposite phenotype when compared to *ric4* mutants.

**Figure 2:**
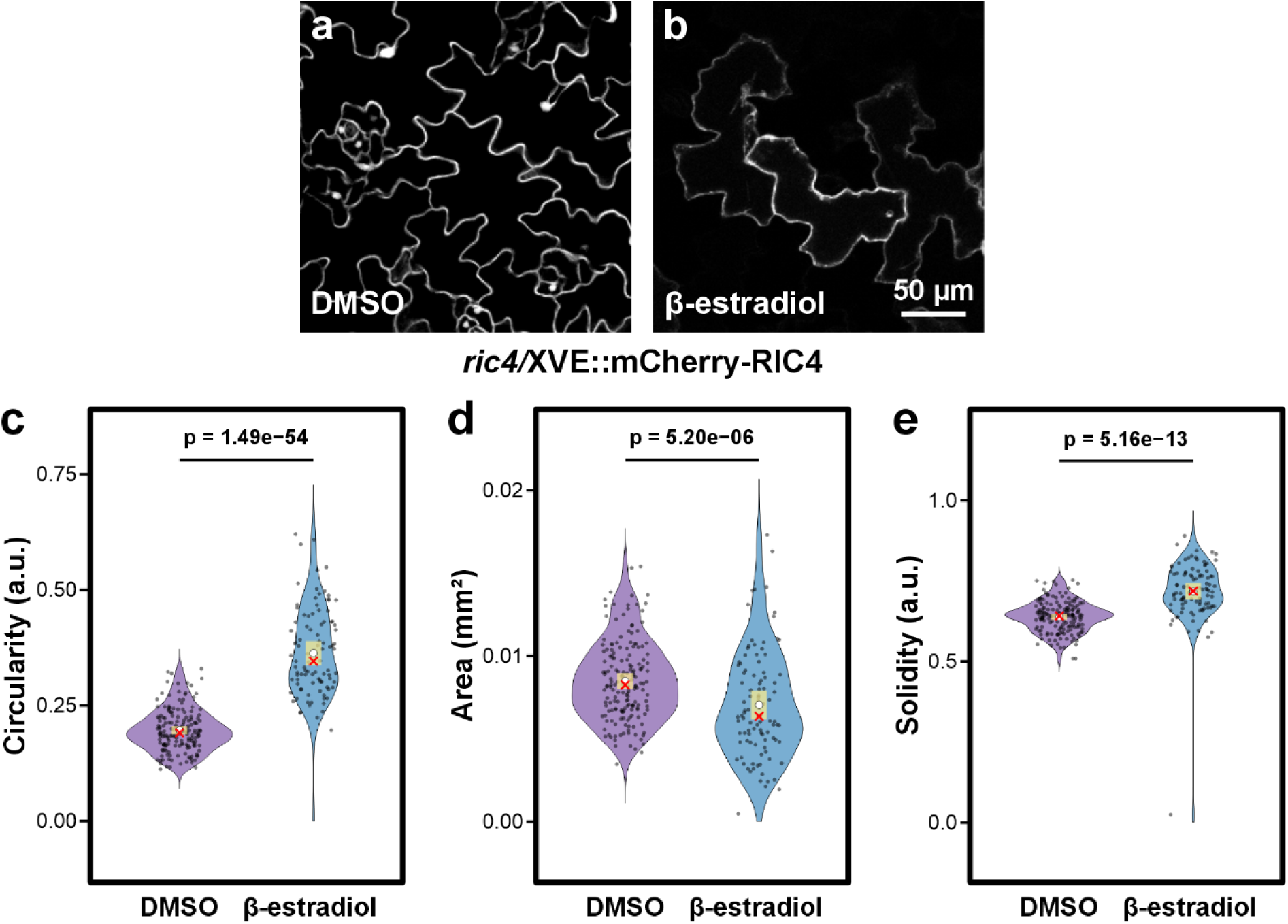
Cotyledon pavement cell analysis of *ric4* mutant expressing mCherry-RIC4 in an estradiol-activated XVE system. (a-b) Representative pictures of propidium iodide-stained cotyledon pavement cells shown as maximum-intensity projections, of uninduced, mock (0.1% DMSO)-treated cells (a) and mCherry-RIC4 expressing plants induced by 2nM β-estradiol (b). (c-e) Violin plots of circularity (c), area (d) and solidity (e) of cotyledon pavement cells for each treatment (total cells: mock, n=172; treated, n = 98 cells across biological replicates). Distributions are overlaid with 99% confidence intervals (yellow bar), mean (white circle) and median (red cross). Statistical comparisons were made using beta-regression GLMMs for circularity and solidity, and a log-transformed linear mixed model for area, followed by Tukey’s HSD-adjusted pairwise contrasts. A significance threshold of α=0.01 (99% confidence) was applied. Scale bar = 50 µm.

We next tested whether overexpression of RIC4 would have an effect in plants lacking a functional Arp2/3 complex. We expressed RIC4 in an inducible vector in Arp2/3 *arpc4* mutants and compared the effect of RIC4 overexpression on cell morphology. RIC4 overexpression clearly induced a significant loss of cell shape complexity in transgenic *arpc4* plants induced with β-estradiol (**Fig. 3d, e, g**), whereas the cell shape was unchanged in mock-treated plants (**Fig. 3c, e, g**). No influence on cell size was detected (**Fig. 3b**). In the same experiment, we confirmed that neither DMSO (**Fig. 3a, e-g**) nor β-estradiol (**Fig. 3b, e-g**) treatment alone had an effect in *arpc4* non-transformed plants (**Fig. 3**). This result demonstrated that the pathway using the RIC4 protein is not dependent on the Arp2/3 complex, because the overexpression of RIC4 effectively influenced the cell shape in the absence of active Arp2/3 complex.

**Figure 3:**
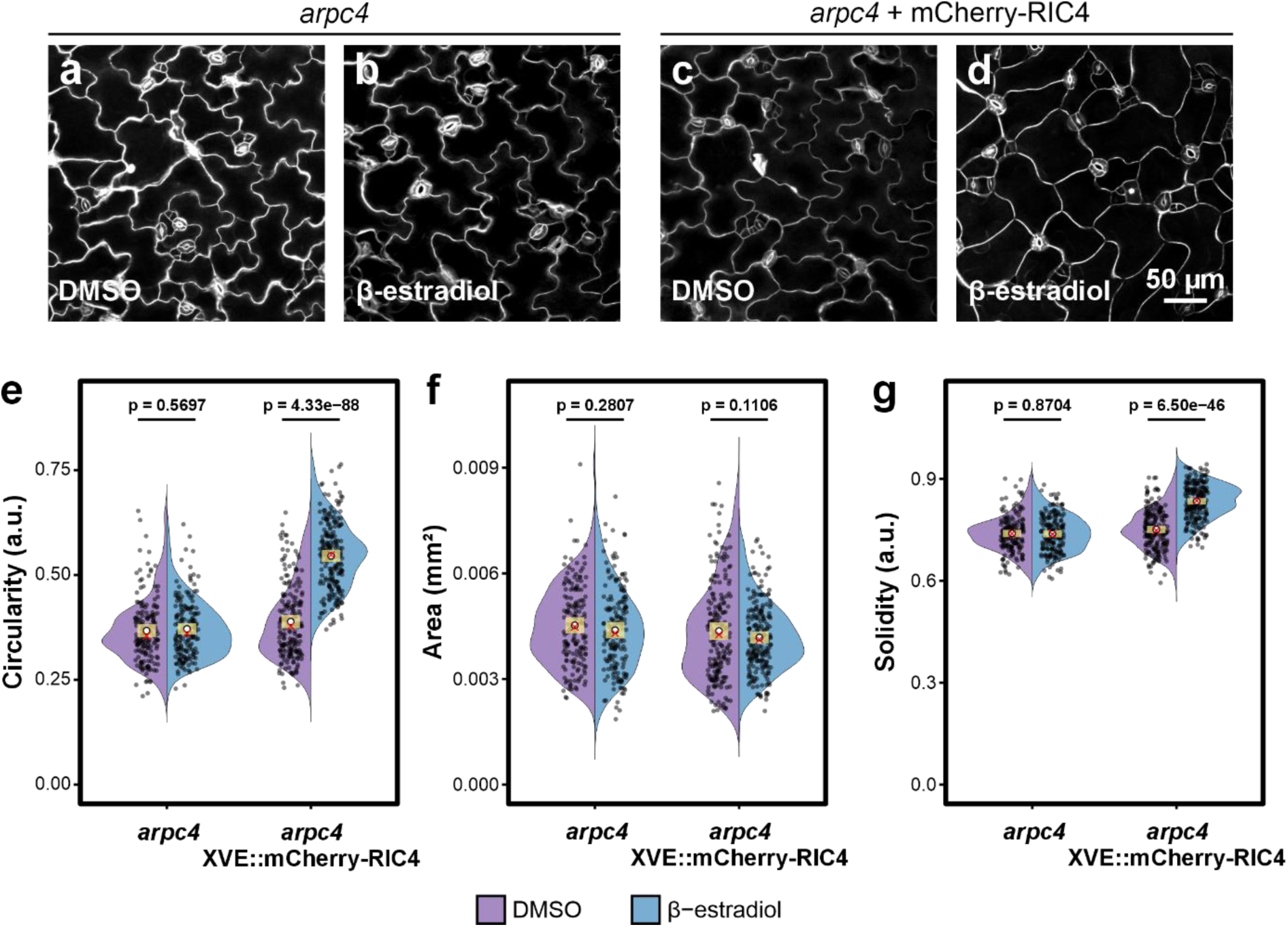
Cotyledon pavement cell analysis of Arp2/3 mutant *arpc4* expressing mCherry-RIC4 induced by the β-estradiol-activated XVE system. (a-d) Maximum-intensity z-projections of propidium iodide-stained cotyledon epidermal cells of uninduced, mock (0.1% DMSO)-treated plants of *arpc4* (a) and *arpc4/XVE::mCherry–AtRIC4* (c) (n=167 and n=201 cells, respectively) and 2nM β-estradiol-treated *arpc4* (b) and *arpc4*/*XVE::mCherry:AtRIC4* (d) (n=176 and n=207, respectively) plants. (e-g) Violin plots of circularity (e), area (f) and solidity for each genotype and treatment combination. Model fitting for was done via beta-regression GLMM for circularity and solidity, and a log-transformed LMM for area with Tukey’s HSD-adjusted pairwise contrasts (α = 0.01). Distributions are overlaid with 99% confidence intervals (yellow bar), mean (white circle) and median (red cross). Scale bar = 50 µm.

We examined whether the same relationship between the Arp2/3 complex and the ROP signaling pathway holds true for the ROP2 protein. *CA-rop2* and *phgap1/phgap2* mutants are defective in ROP2 inactivation, leading to a distinct phenotype of loss of lobes in epidermal cells (Fu et al. 2002; Lauster et al. 2022) (**Supp. Fig 3**). We generated the double and triple mutants of *CA-rop2/arpc5* and *phgap1/phgap2/arpc5* and compared the phenotype of epidermal cotyledon shape. Circularity, solidity, and area were statistically significantly increased in the *arpc5* mutant. However, *CA-rop2* and *phgap1/phgap2* caused a much more dramatic effect that was virtually unaffected by whether the Arp2/3 complex was active in the plants (**Supp. Fig 3**). Thus, both ROP2 gain-of-function and PHGAP1/2 loss-of-function reshape pavement cells via a pathway that bypasses Arp2/3 for lobe control. Arp2/3 loss showed an additive influence on overall cell-surface area only in *CA-rop2* plants but not in *phgap1/phgap2* or RIC4 overexpressing plants. This further suggests that the Arp2/3 complex does not appear to be a primary signaling node downstream of ROP GTPases.

### Cortical actin cytoskeleton has reduced dynamics in cells overexpressing RIC4

Because the function of both ROP2 and RIC4 is associated with the actin cytoskeleton, we explored the relationship between RIC4 and actin in epidermal cells. We crossed mCherry-RIC4 expressing plants with the actin cytoskeleton marker Lifeact-GFP. We analyzed the dynamics of the actin cytoskeleton in RIC4-expressing cotyledon epidermal cells. The method employed correlation coefficient-based analysis that relies on the change in fluorescent pixel intensities over all temporal pairs from a time-lapse series (Vidali et al. 2010).

We compared actin dynamics in the cortical region in induced (RIC4 overexpressing) and uninduced (mock treated) plants. The results clearly showed a significant reduction in actin cytoskeleton dynamics in cells that overexpressed RIC4 protein (**Fig. 4**). To rule out the possibility that actin dynamics changes in induced cells was influenced by the inducing chemical (*β*-estradiol), we used the same method of actin dynamics analysis to compare the dynamics of the actin cytoskeleton in wild-type plants expressing Lifeact-GFP that were cultured on medium with the *β*-estradiol and on medium with the solvent DMSO. The experiments showed that *β*-estradiol- and DMSO-treated plants had comparable cortical actin dynamics (**Supp. Fig. 4a**). Thus, the reduced dynamics in RIC4 overexpressing cells was due to RIC4 overexpression, not to the inducing chemical *β*-estradiol.

**Figure 4:**
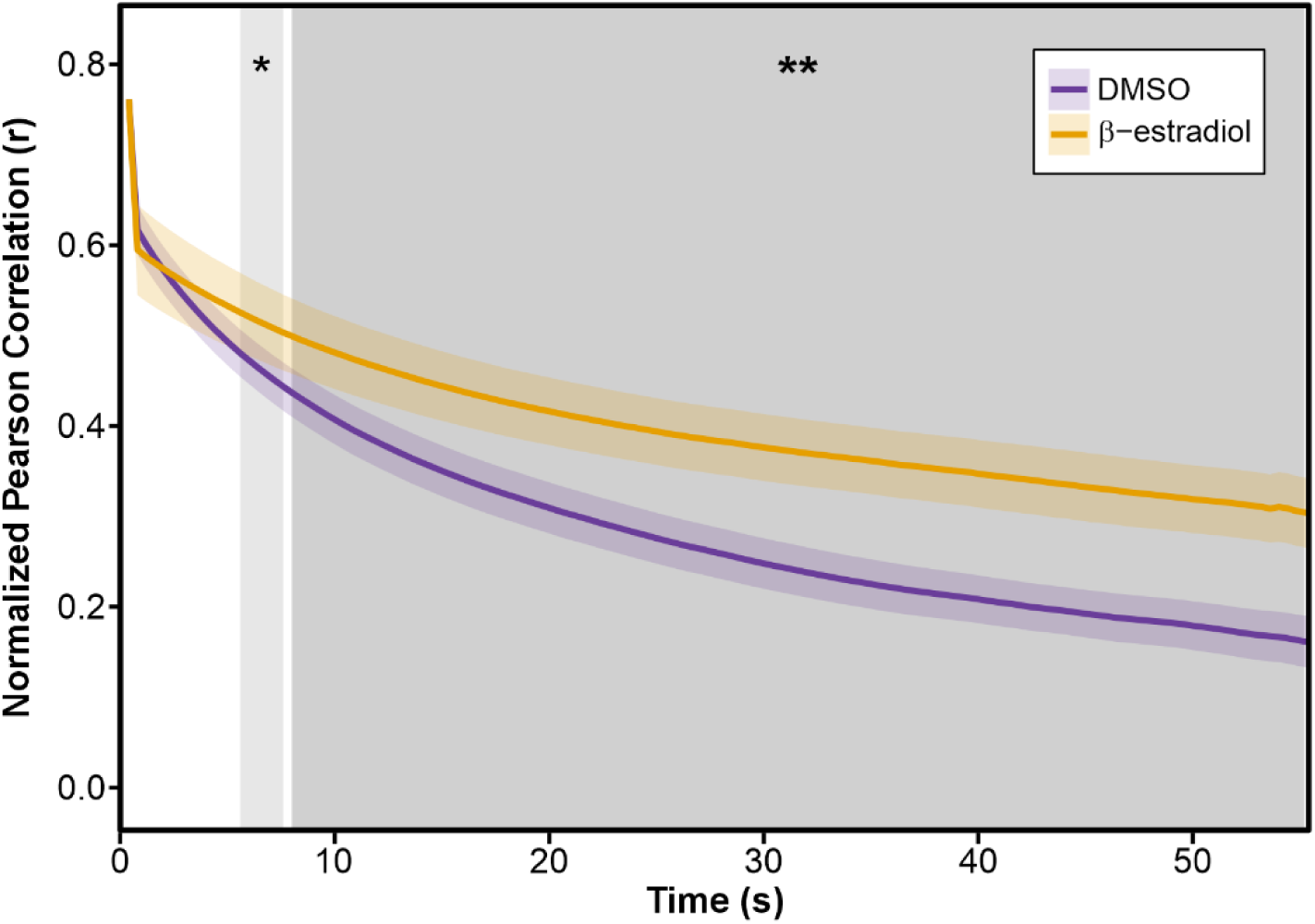
Inducible mCherry–RIC4 expression stabilizes actin filaments in Arabidopsis cotyledon pavement cells. The actin reporter *UBQ::Lifeact-GFP* was introduced into the *ric4*/*XVE::mCherry-AtRIC4* background, and seedlings were mock-treated with 0.1% DMSO (purple) or induced with 2nM β-estradiol (orange). Statistical analysis was performed on Fisher z-transformed correlation coefficients, and data was back-transformed for visualization purposes. Grey shading marks time windows in which β-estradiol-treated cells differ significantly from mock-treated cells (* p < 0.05; ** p < 0.01; two-tailed t-tests with Benjamini-Hochberg correction). Across three independent biological replicates, three cotyledons per treatment—except in the third replicate, where two induced cotyledons were available—were analyzed. Within each plant, three to four regions across the two cotyledons were imaged, and within each region two to four regions of interest (ROIs) were quantified, yielding a total of 53 ROIs for DMSO controls and 47 ROIs for β-estradiol treated plants.

Interestingly, the effect of RIC4 overexpression on actin dynamics was much less dramatic in hypocotyl pavement cells (**Supp. Fig. 4b**). This suggests that the capacity of RIC4 to mediate filaments stabilization depends on the unique morphogenetic context of cotyledons, where specific signaling modules, protein partners, and cell-shape cues converge to sensitize the cortex to RIC4’s action.

### The relationship between cortical actin dynamics and cell shape changes is inconclusive

RIC4 overexpressing cells have increased circularity, suggesting a negative effect of RIC4 on lobes formation. RIC4 overexpressing cells also have reduced cortical actin network dynamics. Mutants without functional Arp2/3 complex develop epidermal cells of increased circularity as well. To determine whether our method of actin dynamics analysis could demonstrate a relationship between cortical actin dynamics and the process of pavement cells shape formation, we compared the dynamics of wild-type plants and *arpc5* plants cortical actin (**Fig. 5**). Therefore, we crossed an *arpc5*/*UBQ10::Lifeact-GFP* plant to wild-type Col-0 plant to generate *arpc5*^⁺/⁻^ *UBQ10::Lifeact-GFP*^⁺/⁻^ progeny, and in parallel to an *arpc5* homozygote to obtain *arpc5*⁻/⁻ *UBQ10::Lifeact-GFP*^⁺/⁻^ F1 plants. Because both lines were derived from the same Lifeact-GFP–expressing parent, they exhibited comparable levels of marker fluorescence. This ensured that any observed differences in cortical actin dynamics arose from the Arp2/3 genotype rather than from variation in Lifeact-GFP expression, which is known to bias cytoskeletal measurements (Cvrčková and Oulehlová 2017). In our previous research, we showed that *arpc5* plants, which have increased circularity of pavement cells in cotyledons, have slightly increased bundling (skewness) and occupancy, but the dynamic properties measured as lifetime of actin using QuACK method (Cvrčková and Oulehlová 2017) were not changed (Cifrová et al. 2020). In this work, we have combined spinning disc confocal time series imaging with the correlation coefficient-based method for actin dynamics measurements. Our results demonstrate that *arpc5*^+/-^ and *arpc5*^-/-^ plants show similar dynamics of actin (**Fig. 5**), confirming our previous results. Although RIC4 overexpression in cotyledons markedly stabilizes cortical actin and reduces the cell shape complexity, the absence of any detectable decrease in overall actin dynamics in *arpc5* mutants—despite their similar high circularity and solidity—indicates that bulk actin-dynamics measurements can miss local, lobe-specific changes in filaments behavior. Consequently, reduced actin dynamics and increased circularity cannot be assumed to be causally linked based on whole-cell assays alone.

**Figure 5:**
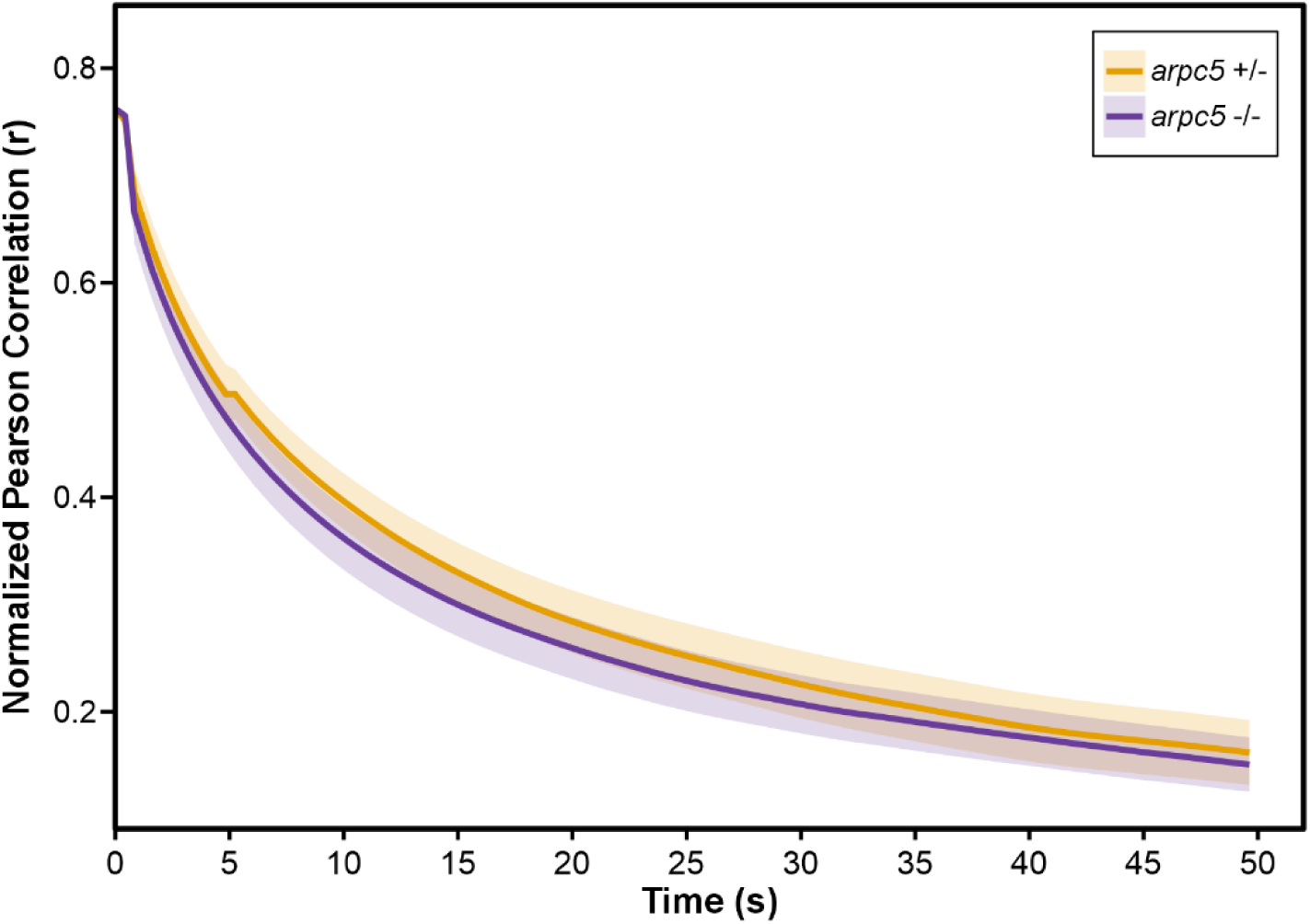
Correlation coefficient analysis of actin dynamics in *arpc5* ^+/-^ and *arpc5 ^-/-^* plants. Actin marker *UBQ10::Lifeact-GFP* was introduced into *arpc5* mutants and crossed with wild-type and *arpc5* mutants. The resulting F1 generation of heterozygous (*arpc5* ^+/-^) and homozygous (*arpc5* ^-/-^) plants expressing Lifeact-GFP were used for actin dynamics evaluation in cotyledon pavement cells imaged using spinning-disc confocal microscope. Correlation coefficients (normalized Fisher z-transformed, back-transformed for visualization) showed no significant difference between *arpc5* ^+/-^ and *arpc5* ^-/-^ (two-tailed t-tests with Benjamini–Hochberg correction, α=0.05). Data were collected across three independent biological replicates: in replicate 1, three *arpc5* ^+/-^ and three *arpc5* ^-/-^ plants; in replicate 2, two plants of each genotype; and in replicate 3, three *arpc5* ^+/-^ and four *arpc5* ^-/-^ plants were imaged. Within each plant, two to four distinct cotyledon regions were imaged, and 1–5 regions of interest (ROIs) per imaged area were quantified—total 56 ROIs for *arpc5* ^+/-^ and 62 ROIs for *arpc5* ^-/-^.

### RIC4 and actin are not enriched in cell lobes, but RIC4 overexpressing cells show mild actin structure changes

We further attempted to establish a link between the organization of the actin cytoskeleton and the localization of the RIC4 protein. The motivation for this experiment came from studies in which fine actin was detected in developing lobes, and its abundance was further increased in cells overexpressing RIC4 (Fu et al. 2005). In our experimental material, mCherry-RIC4 was evenly distributed in the cytoplasm in cells overexpressing RIC4 and no specific enrichment was observed in lobes or other cell compartments (**Fig. 6a, b**). Small round motile organelles of unknown origin that were positive for RIC4 protein were also regularly detected in the cells. These structures were also uniformly distributed in the cell and their preferential localization in lobes was not observed (**Fig. 6b**). The uniform distribution of the protein was particularly evident, where RIC4 was expressed as a mosaic in few epidermal cells, which allowed precise 3D microscopical analysis and reconstruction. The actin cytoskeleton in cells overexpressing mCherry-RIC4 did not show increased depolymerization or enrichment in cell lobes (**Fig. 6c, d**). For quantitative evaluation of actin structure in cells overexpressing mCherry-RIC4, we analyzed actin in the cortical layer of epidermal cells, where we previously observed reduced actin dynamics. The reason for this is that current methods of actin analysis do not allow for accurate analysis of actin structure directly in lobes. We used several well-known parameters to characterize actin organization: the coefficient of variation (cv) and skewness to quantify the degree of bundling (Higaki et al. 2020), anisotropy to measure directionality of the actin network, and occupancy as a measure of actin density. First, we evaluated actin structure in wild-type cells expressing the actin marker, treated with DMSO (mock) and β-estradiol. We found that there was no difference in any of the parameters between wild-type cells treated with DMSO and β-estradiol (**Fig. 6e-h**), confirming that β-estradiol itself does not affect the cells. Analysis of the actin structure in cells overexpressing mCherry-RIC4 showed a reduced cv factor at a low level of significance compared to cells in which overexpression was not induced (**Fig. 6e**). This suggests that actin in cells with induced expression of mCherry-RIC4 tends to form less bundled actin. Skewness, another widely used parameter to quantify filament bundling, also confirmed the observation (**Fig. 6g**). We therefore concluded that mCherry-RIC4 expression induces a slight reduction of actin bundling. The occupancy factor showed a slight increase in induced cells, also at a very low level of statistical significance (**Fig. 6h**). Anisotropy quantifies the degree to which actin filaments share a common orientation—high anisotropy indicates strongly aligned bundles, whereas low anisotropy reflects a more random filament network. In cells expressing mCherry-RIC4, reduced anisotropy was detected (**Fig. 6f**). Our results showed that induction of mCherry-RIC4 overexpression in cotyledon epidermal cells results in rather minor structural changes in the sense of forming less bundled actin, increasing actin network density, and a tendency to form more randomly organized arrays, which could affect the polarity of actin-based processes that are necessary for proper shape formation.

**Figure 6:**
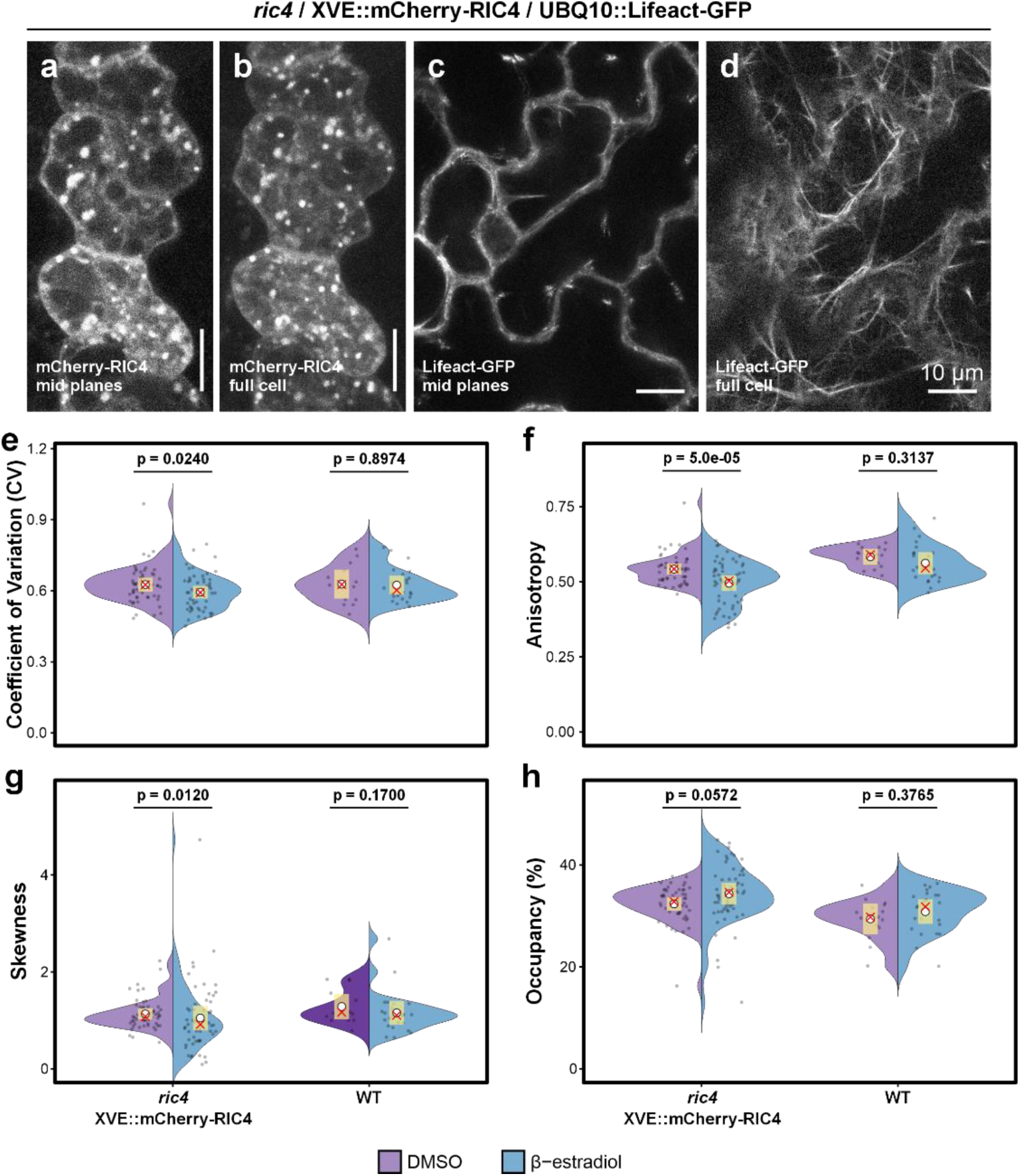
Cortical actin organization in pavement cells expressing mCherry–RIC4 versus wild-type controls. (a-b) mCherry-RIC4 in pavement cells is distributed homogenously in the cytoplasm and in small organelles. Maximum intensity projection of four central optical sections (a); maximum intensity projection of optical sections through whole cell thickness (b). (c-d) Organization of actin cytoskeleton in cells expressing Lifeact-GFP shows no specific accumulation in lobes. Maximum projection of five central optical sections (c); single optical section through the cortical cytoplasm (d). Confocal microscopy, scale bar = 10 µm. (e-h) Quantification of cortical actin organization in *ric4*/*XVE::mCherry-AtRIC4*/*UBQ10::Lifeact-GFP* versus wt/*UBQ10::Lifeact-GFP* under mock (0.1% DMSO, purple) or 2nM β-estradiol (blue) induction. Shown are coefficient of variation (e), anisotropy (f), skewness (g), and occupancy (h). Data derive from three independent biological replicates of the *ric4/XVE::mCherry-AtRIC4* line, three plants per treatment per replicate; 2–8 cotyledon regions per plant; 1–3 ROIs per region), yielding 52 ROIs in the DMSO control and 55 ROIs under β-estradiol. wt controls were processed in parallel (three plants per treatment; 1–6 regions per plant; 2–3 ROIs per region), for 15 ROIs (DMSO) and 27 ROIs (β-estradiol). In the violin plots, the yellow bar denotes the 99% confidence interval, the white circle marks the mean, and the red cross indicates the median. Occupancy and anisotropy were analyzed by beta-GLMM (logit link, estimated marginal means contrasts, Tukey adjustment), skewness by robust linear regression on the log-transformed values with bootstrapped contrasts (2000 iterations), and CV by linear regression on log-transformed values (estimated marginal mean contrasts, Tukey adjustments). For all statistical analysis, two-tailed tests were performed, and p-values were adjusted for multiple comparisons (Benjamini-Hochberg).

## Discussion

The aim of this study was to determine whether RIC4 and Arp2/3 complex are functionally related in the control of Arabidopsis epidermal cell shape. Both factors are linked to the actin cytoskeleton and the formation of specific puzzle-like epidermal cell shapes. Analysis of phenotypes of single and double mutants showed that Arp2/3 and RIC4 have distinct roles. Loss of the Arp2/3 complex (*arpc5*) resulted in increased cell circularity, consistent with previous studies (Pratap Sahi et al. 2017; Cifrová et al. 2020). However, careful cell shape analysis showed that single KO mutant *ric4* has decreased circularity of cotyledon pavement cells, which suggests that the cells have increased lobes formation and cell shape complexity. Moreover, the *arpc5/ric4* double mutant clearly showed principally the same morphological phenotype as the *arpc5* single mutant, although mild additive effects were also present. Generally, loss of the Arp2/3 complex masked the effect of the *ric4* mutation. This result ruled out the possibility that RIC4 affects the actin cytoskeleton through activation of Arp2/3. Interestingly, unchanged pavement cell shape of *ric4* mutant was reported by (Belteton et al. 2018). The discrepancy of the two analyses is probably related to different developmental stages of analyzed plants and cultivation conditions, where our study used fully expanded cotyledons of 14-day-old plants cultivated at 8/16 photoperiod.

Both of studied factors are involved in the control of cell shape by the actin cytoskeleton, but the molecular mechanism of this process is unknown. Arp2/3 is a conserved actin nucleator and it is proposed that actin nucleated by Arp2/3 is specifically involved in epidermal cell expansion and lobes formation. RIC4 is not a direct actin-binding protein, nor is there any other known mechanism of interaction with actin, but its overexpression has been proposed to induce the formation of fine actin in epidermal cells at the site of the lobes (Fu et al. 2005). However, since we have shown that loss of *ric4* results in increased formation of complex cell shapes, it is clear that the function of RIC4 is in fact lobes formation inhibition. This hypothesis is supported by our further observation that overexpression of mCherry-RIC4 in cotyledon epidermal cells dramatically increases circularity. Importantly, these RIC4-induced cell shape changes occurred also in mutants lacking functional Arp2/3 (*arpc4*) (Pratap Sahi et al. 2017). These observations have ruled out the possibility that Arp2/3 and RIC4 are functionally dependent components of the same signaling pathway controlling cell shape formation.

RIC4 is a putative downstream effector of ROP GTPases (Wu et al. 2001), which are involved in the regulation of epidermal cell shape (Yang 2002). It is probable that the overexpression phenotype is the result of an imbalance of ROP signaling in epidermal cells rather than a specific RIC4 protein function. Imbalance of ROP signaling components usually leads to similar dramatic phenotypes (Kost et al. 1999; Li et al. 1999; Fu et al. 2002). Therefore, we compared the effect of *CA-rop2* expression (Fu et al. 2002) and the effect of the *phgap1/phgap2* mutation (Lauster et al. 2022) in plants lacking a functional Arp2/3 complex (*arpc5* and *arp2*). We found that the effect of *CA-rop2* and *phgap1/phgap2* is consistently independent of the existence of an active complex, although small additive effects could be measured in cell shape and size traits also here. We concluded that the Arp2/3 complex is not a primary downstream effector of RIC4 protein or general ROP signaling in epidermal cells. However, mild additive effects in double mutants *arpc5/ric4*, *arpc5/CA-rop2* and *arp2* or *arpc5* combined with *phgap1/phgap2* triple mutants also leads us to a suggestion that Arp2/3 and ROP signaling functions rather in parallel pathways and we cannot exclude the possibility of their cooperation in pavement cell morphogenesis.

We attempted to determine the correlation between the phenotype of overexpressed RIC4 and the actin cytoskeleton. According to our analysis of the structure and distribution of the actin cytoskeleton, there is no specific actin remodeling in the lobes or elsewhere in the cells of plants expressing Lifeact-GFP and mCherry-RIC4. We did not see any fine actin localized specifically in lobes. To learn as much as possible about the actin cytoskeleton in these cells, we focused on actin dynamics and actin structure in the cortical layer of epidermal cells. Because actin dynamics in lobes is difficult to measure with current microscopic methods, we focused on the cortical actin in epidermal cells at the outer periclinal wall and we used a method based on correlation coefficient measurements. We hypothesized that if RIC4 influences shape changes through the actin cytoskeleton, we should detect a change in actin in overexpressing cells. The results clearly showed that cells overexpressing RIC4 have a more stable actin cytoskeleton. Surprisingly, the analysis of actin structure showed only mild changes induced by mCherry-RIC4, which were slightly decreased bundling, increased density and decreased anisotropy of actin arrays, all detected at a low level of statistical significance with the exception of statistically more significant changes in anisotropy.

If simple actin stabilization measured with our approach in the cortical region of epidermal cells is the main factor responsible for the increase in cell circularity, we should also detect actin stabilization in cells lacking the Arp2/3 complex, whose phenotype is also increased circularity. However, correlation coefficient analysis showed no significant changes in actin dynamics in Arp2/3 mutants. Thus, while our results confirmed a functional relationship between RIC4 and actin, they also suggested that a clear functional link between changes in actin dynamics and changes in cell shape could not be established. Actin dynamics and cell shape changes may be completely independent phenomena. Interestingly, we were unable to demonstrate similarly dramatic change in actin dynamics in hypocotyl cells that overexpressed mCherry-RIC4. This means that actin structure and regulation mechanisms are rather tissue or developmentally specific in Arabidopsis seedling. Importantly, cotyledon epidermal cells must possess a regulation pathway that responds to mCherry-RIC4 expression in contrast to hypocotyl epidermal cells. In future studies, this finding could lead to the identification of a cellular factor required for the function of RIC4.

This study certainly has limitations in the context of actin analysis. We measured actin dynamics just below the plasma membrane on the periclinal walls of cotyledon epidermal cells, but not actin dynamics in the lobes themselves, where this is technically impossible because of the high abundance and dynamics of actin and limited visibility by microscopes. In our experiments, we were aware of the influence of marker expression on actin dynamics (Cvrčková and Oulehlová 2017), so we carefully compared only marker lines derived from crosses between a single marker line (*arpc5*^-/-^ vs *arpc5*^+/-^). In the case of RIC4 overexpressing lines, we used inducible expression to compare actin dynamics in cells of a single line. Finally, we verified that induction by β-estradiol alone does not affect actin dynamics. However, the actin cytoskeleton is an abundant structure, and it is very likely that the function of Arp2/3-nucleated actin is highly localized in the cell and therefore does not affect the overall dynamics of the actin network. This is suggested by our previous results, when we were able to demonstrate very little change in the overall actin cytoskeleton dynamics of the cotyledon epidermal cells of Arp2/3 mutant *arpc5* in confocal microscope images (Cifrová et al. 2020) using semi-manual method Quantitative Analysis of Cytoskeletal Kymograms (QuACK) that evaluates kymograms constructed based on manually positioned transects (Cvrčková and Oulehlová 2017). In the work of (Xu et al. 2024) in etiolated hypocotyls, detailed analysis did reveal changes of the actin structure and dynamics in mutants lacking the Arp2/3 complex. The mutants had reduced filament abundance and bundling. At the single filament level, reduced frequency of lateral nucleation, longer filament length and longer lifetime were detected (Xu et al. 2024). Since our present results suggest that actin in different organs shows different responsiveness to regulatory factors, it is probable that the data from dark-grown hypocotyls and expanded light-grown cotyledon pavement cells cannot be compared. Our analysis suggests that Arp2/3 mutants do not have affected overall actin dynamics in cotyledons when compared to wild-type, and structural analysis revealed only minor changes – slightly increased density, decreased bundling and decreased anisotropy in cotyledon pavement cells. Yet the same cells have a phenotype of increased circularity when Arp2/3 complex is not available. Therefore, our present results suggest that to understand the role of actin in specific cellular processes, we need to study the structure and dynamics of actin at these specific sites, because the general structure or dynamics of abundant actin in other regions does not need to be affected by local processes. In many cases, our current technical capabilities are insufficient for such observations.

## Acknowledgement

Confocal and TIRF microscopy was performed in the Vinicna Microscopy Core Facility (RRID:SCR_026602) co-financed by the Czech-BioImaging large RI project LM2023050. Computational resources were supplied by the e-INFRA CZ project (ID:90254) provided within the program Projects of Large Research, Development, and Innovations Infrastructures. Spinning disc microscopy was performed at the Imaging Facility of the Institute of Experimental Botany AS CR supported by the MEYS CR (LM2023050 Czech-Bioimaging), the Czech Academy of Sciences and IEB AS CR.

## Author contribution

KS and JG conceived and designed research. JG, LH, EB, IC, PP, MP and KS conducted experiments. JG and KS analyzed data. KS and JG wrote the manuscript. All authors read, corrected and approved the manuscript.

