## Supplementary data 1 and 2 for "Independent roles of Arp2/3 complex and RIC4 protein in the control of epidermal cell shape"

**Statistical analysis of actin dynamics:**

To enable statistical comparisons across ROIs with non-uniform time grids, Fisher z-transformed correlation values were linearly interpolated onto a common time grid. The interpolation step used the median observed time interval and was restricted to the measured time range of each ROI (i.e., no extrapolation). Interpolated z-scores were normalized within each ROI by dividing each value the value at the initial timepoint, then back-transformed to correlation values (r) using the hyperbolic tangent function for later visualization.

To assess dynamic treatment or genotype effects over time, a generalized linear mixed model was fitted using the glmmTMB package. The response variable was the normalized Fisher z-correlation for each ROI and time point. Time was represented using orthogonal polynomial terms (linear, quadratic, cubic) to capture nonlinear trends. The fixed-effect structure included treatment (or genotype) and the time terms. Biological repetition was included as a random intercept, and ROI identity nested within repetition was modelled with a random slope on time.

The model is given by the following equation:

$$Y_{ijkt} = \beta_0 + \beta_1 \cdot \mathrm{Group}_j + \beta_2 \cdot T1_t + \beta_3 \cdot T2_t + \beta_4 \cdot T3_t + b_{0k} + b_{1ik} \cdot T1_t + \varepsilon_{ijkt}$$

where $\mathrm{Group}_j$ stands for either treatment or genotoype, $T1_t$ ,$T2_t$ ,$T3_t$ are the orthogonal polynomial time terms (linear, quadratic, cubic), $b_{0k}$ is the random intercept for biological repetition k, $b_{1ik}$ is the random slope of time (T1) for ROI i within biological repetition k, and $\varepsilon_{ijkt}$ is the residual error.

Model residuals were checked for temporal autocorrelation using autocorrelation function (ACF) plots per ROI. To address within-ROI autocorrelation, an ARIMA(1,1,0) correction was applied individually per ROI on the model residuals. The resulting fitted ARIMA trends were added to the initial mixed-model predictions to produce ARIMA-adjusted predicted values, which were used for further statistical inference.

An updated mixed model was then fit incorporating the ARIMA correction as an offset term, allowing standard inference while preserving autocorrelation structure in the data. The model was defined as:

$$Y_{ijkt} = \beta_0 + \beta_1 \cdot \mathrm{Group}_j + \beta_2 \cdot T1_t + \beta_3 \cdot T2_t + \beta_4 \cdot T3_t + b_{0k} + b_{1ik} \cdot T1_t + \varepsilon_{ijkt} + offset_{ijkt}$$

where all terms correspond to the original fitted model, with an additional $\mathrm{offset}_{ijkt}$ term, corresponding to the ARIMA correction applied to each time point.

Residual diagnostics were repeated on the offset model, and model comparison was conducted using AIC. Additional model checks were performed to check for validity of assumptions, namely residual QQ plots, Saphiro-Wilk tests and checks for homoscedasticity and dispersion.

To evaluate treatment or genotype effects over time, pairwise t-tests were conducted on ARIMA-corrected predicted z-values at each time point. P-values were adjusted for multiple comparisons using the Benjamini-Hochberg method. Significant contrasts between treatments or genotypes were visualized over time, highlighting significant windows with shaded regions based on adjusted significance level.

### Supplementary Data 2:

**Image enhancement and segmentation for actin structure analysis:**

To enhance the visibility of filamentous structures while minimizing background interference, we employed the Implicit Laplacian of Enhanced Edge (ILEE, v1.0) method, originally described by Li and colleagues (Li et al. 2023) and adapted from the open-source implementation (available at https://github.com/phylars/ILEE_CSK). Briefly, ILEE is a local edge-based filtering algorithm that detects filament edges and solves a Poisson equation to reconstruct an intensity surface that conforms to local gradients. This adaptive contrast enhancement avoids global thresholding and is particularly suited to cytoskeletal networks with spatially variable signal.

ILEE filtering was applied independently to each time point in the 2D+t image stacks. The enhanced image sequences were stored and served as input for downstream filament detection.

To further highlight curvilinear filament structures and suppress isotropic noise, we applied a multiscale tubeness filter (Frangi filter) to the ILEE-enhanced images using the python library scikit-image (0.24.0) implementation. This tubeness filter enhances regions in the image with high likelihood of being part of curvilinear structures (based on the eigenvalues of the Hessian matrix), suppressing isotropic features and noise. This step produces smooth, filament-specific intensity maps that are well suited for deep learning-based segmentation.

Segmentation of actin filaments was performed using a Densely Connected Stacked U-Net (DSC-UNet), adapted from the model published by Liu and colleagues (Liu et al. 2018, 2019) (available at https://github.com/i-yliu/mtquant). This architecture consists of multiple U-Net blocks connected in series with dense residual connections, which improves segmentation performance on thin and faint filament structures by propagating multi-scale features through the network.

In our pipeline, the pretrained model available for actin filament segmentation available directly from the repository (weight_4_1_2020.h5) was used in inference mode to segment the actin cytoskeleton in each frame of the 2D+t sequence independently. Prediction was performed in a sliding-window fashion using 64×64 pixel patches with batch size 8. The resulting probability maps were thresholded to generate binary masks of filament regions. All segmented masks were saved as .tif stacks and passed on for structural quantification.

Preprocessed binary segmentation masks—generated via deep learning-based inference on ILEE-enhanced images—were used as the basis for structural and intensity-based analysis. Morphological quantification was carried out on the binary masks and their skeletonized forms, while intensity-based features were derived from the original or enhanced images constrained to the actin-positive regions defined by the masks. Skeletons were further interpreted as graph structures, enabling extraction of topological descriptors.

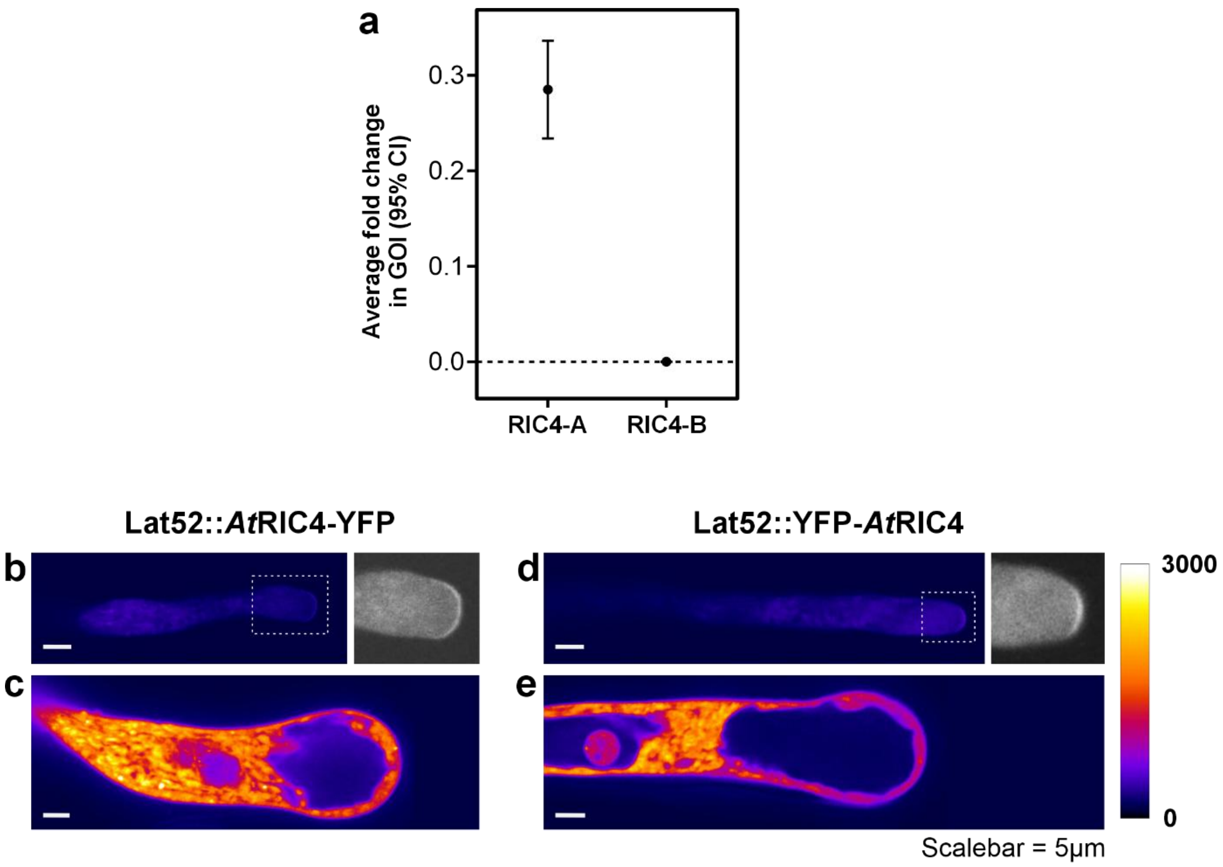


**Supplementary Figure 1**: Validation of the *ric4* T-DNA insertional mutant and functional assay of RIC4 fusions with YFP in tobacco pollen tubes. (a) qRT-PCR of AtRIC4 transcript levels in wild-type versus *ric4* T-DNA insertion line homozygotes. The amplicon upstream of the T-DNA insertion (RIC4-A) is reduced to ~0.3-fold of wild-type, whereas the downstream amplicon spanning the insertion site (RIC4-B) is undetectable. Bars show mean ± 95% Confidence Interval (CI) (n = 3 independent biological replicates, each with three technical replicates). (b–e) Transient expression of *Lat52::AtRIC4:YFP* and *Lat52::YFP:AtRIC4* fusions in *Nicotiana tabacum* pollen tubes. Both C-terminal (RIC4–YFP; b,c) and N-terminal (YFP–RIC4; d,e) fusions concentrate at the tip of pollen tubes. Insets in (b,d) emphasize membrane-associated fluorescence and subapical clustering. At moderate expression levels (b,d), tubes elongate with normal polarity; at higher expression (c,e), they display apical swelling and overall enlargement, confirming that the YFP fusions are functional and perturb the ROP signalling cascade. Scale bars = 5 µm.

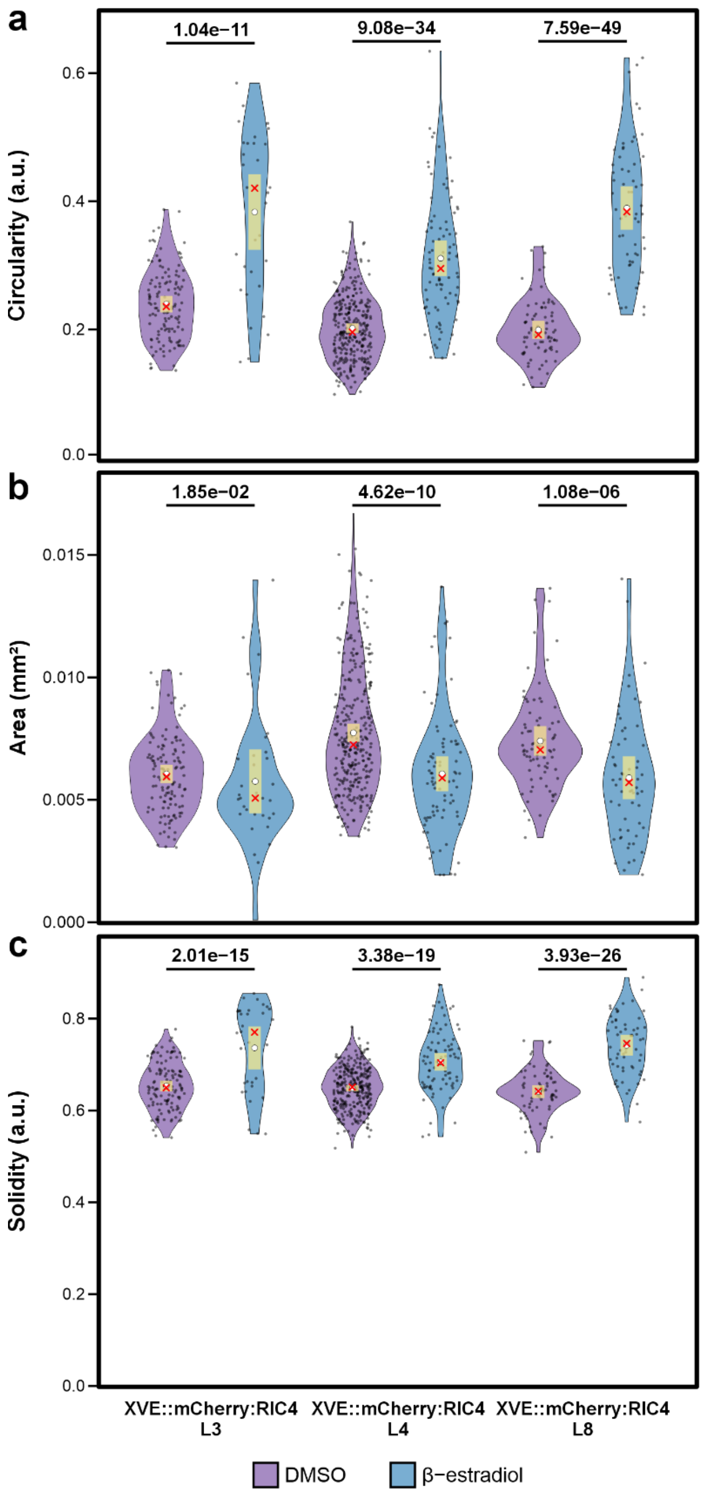


**Supplementary Figure 2**: Analysis the effect of mCherry-RIC4 expression on cell shape in three independent transformants of *ric4* mutants. All three analysed Arabidopsis lines showed increased circularity (a), decreased area (b) and increased solidity (c) parameters of cotyledon pavement cell shape under induction conditions (treatment with β-estradiol) when compared to mock (DMSO) treatment. Across biological replicates, for transformant L3, 126 DMSO-treated and 33 β-estradiol-treated cells were analysed; for transformant L4, 304 and 83 cells; and for transformant L8, 77 and 60 cells, respectively. Violin plots show a yellow bar indicating the 99 % confidence interval around the mean (white circle), and the red cross marks the median. Statistical significance was determined by linear mixed-effects models, followed by pairwise estimated marginal means contrasts with Tukey adjustment for multiple comparisons. P-values are indicated above violin plots.

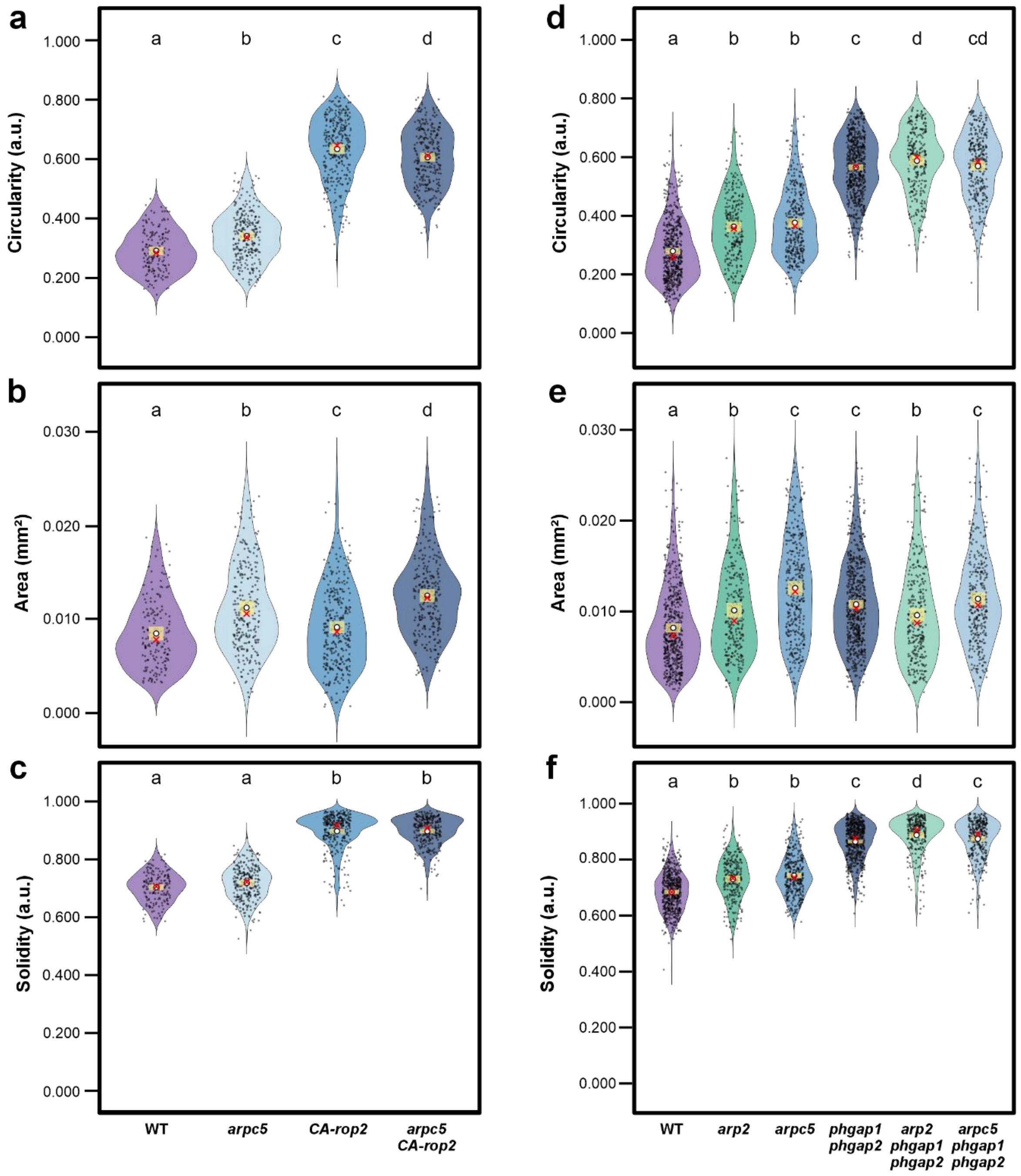


**Supplementary figure 3**: Violin plots of cotyledon pavement cell shape parameters in ROP2 pathway and Arp2/3 complex mutants. Loss of ROP2 inactivation (*CA-rop2* or *phgap1/phgap2*) causes a dramatic increase in each shape parameter that is not further exacerbated by *arpc5* mutation. Panels (a–c) compare wild-type (n = 169 cells), *arpc5* (n = 269), *CA-rop2* (n = 328) and *CA-rop2/arpc5* double mutants (n = 293), pooled from at least three independent biological replicates for (a) cell area, (b) circularity and (c) solidity; panels (d–f) compare wild-type (n = 560), *arp2* (n = 259), *arpc5* (n = 370), *phgap1/phgap2* (n = 643) and the corresponding *arp2/phgap1/phgap2* (n = 294) and *arpc5/phgap1/phgap2* triple mutants (n = 334), pooled from at least three biological replicates for (d) area, (e) circularity and (f) solidity. Violin shapes depict the full distribution of individual cell measurements (points), the yellow bar indicates the 99 % confidence interval of the mean (white circle) and the red cross marks the median. Cell area was analysed on the log scale using a two-level Gaussian mixed-effects model; circularity and Smithson–Verkuilen-adjusted solidity were analysed by β-regression. Pairwise comparisons were performed with Sidak-adjusted multiple-comparison tests (α = 0.01), and groups that differ significantly are denoted by distinct letters above each violin.

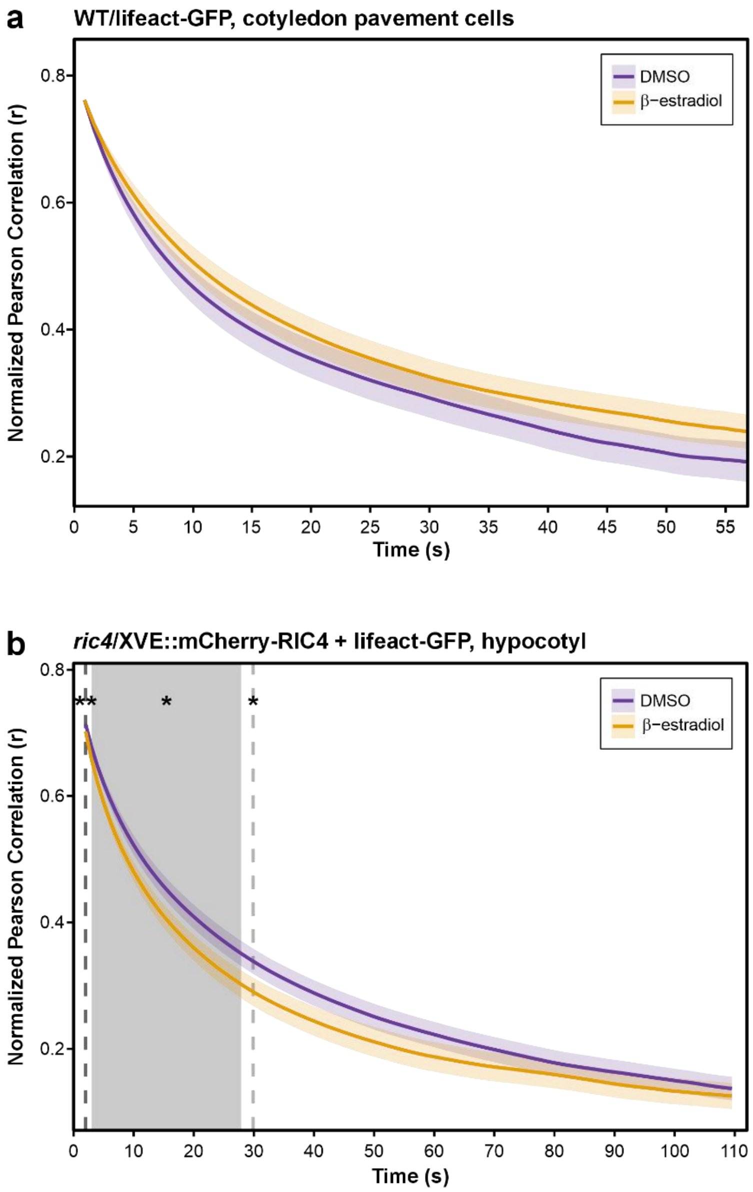


**Supplementary Figure 4**: Correlation coefficient (CV) analysis of actin filament dynamics in (a) wild-type cotyledon pavement cells expressing Lifeact-GFP, and (b) in hypocotyl pavement cells of mCherry-RIC4 induced *ric4* mutant-expressing Lifeact-GFP, imaged by spinning disc confocal microscope. (a) Seedling cotyledons were mock-treated with 0.1% DMSO or induced with 2 nM β-estradiol. Data derive from two independent biological replicates; in each replicate, three plants per treatment were imaged. DMSO-treated plants were sampled in 2–3 distinct cotyledon regions per plant, whereas β-estradiol–treated plants were sampled in 1–5 regions per plant. Within each region, 2–4 regions of interest (ROIs) were quantified, yielding a total of 44 ROIs for DMSO mock controls and 55 ROIs for β-estradiol–treated plants. (b) 5-day-old hypocotyl epidermal cells from *ric4/XVE::mCherry-AtRIC4* expressing Lifeact-GFP were also analysed after β-estradiol-induction of mCherry-RIC4 expression compared to mock-treated. Images were acquired across three biological replicates with at least 4 plants per treatment. For each plant, one to three regions were sampled, and one to five ROIs were selected for each region, yielding a total of 77 ROIs for DMSO controls and 47 ROIs for β-estradiol-treated plants. Normalized Pearson correlation coefficients (tanh-back-transformed from Fisher z) are plotted over time; shaded areas show ± 95 % CI. Grey shading marks time windows in which β-estradiol-treated plant actin dynamics differs significantly from DMSO-treated plants (* $p < 0.05$; ** $p < 0.01$; two-tailed t-tests with Benjamini–Hochberg correction).
