## Supplementary figures and images for "Independent roles of Arp2/3 complex and RIC4 protein in the control of epidermal cell shape"

### Supplementary figure 1

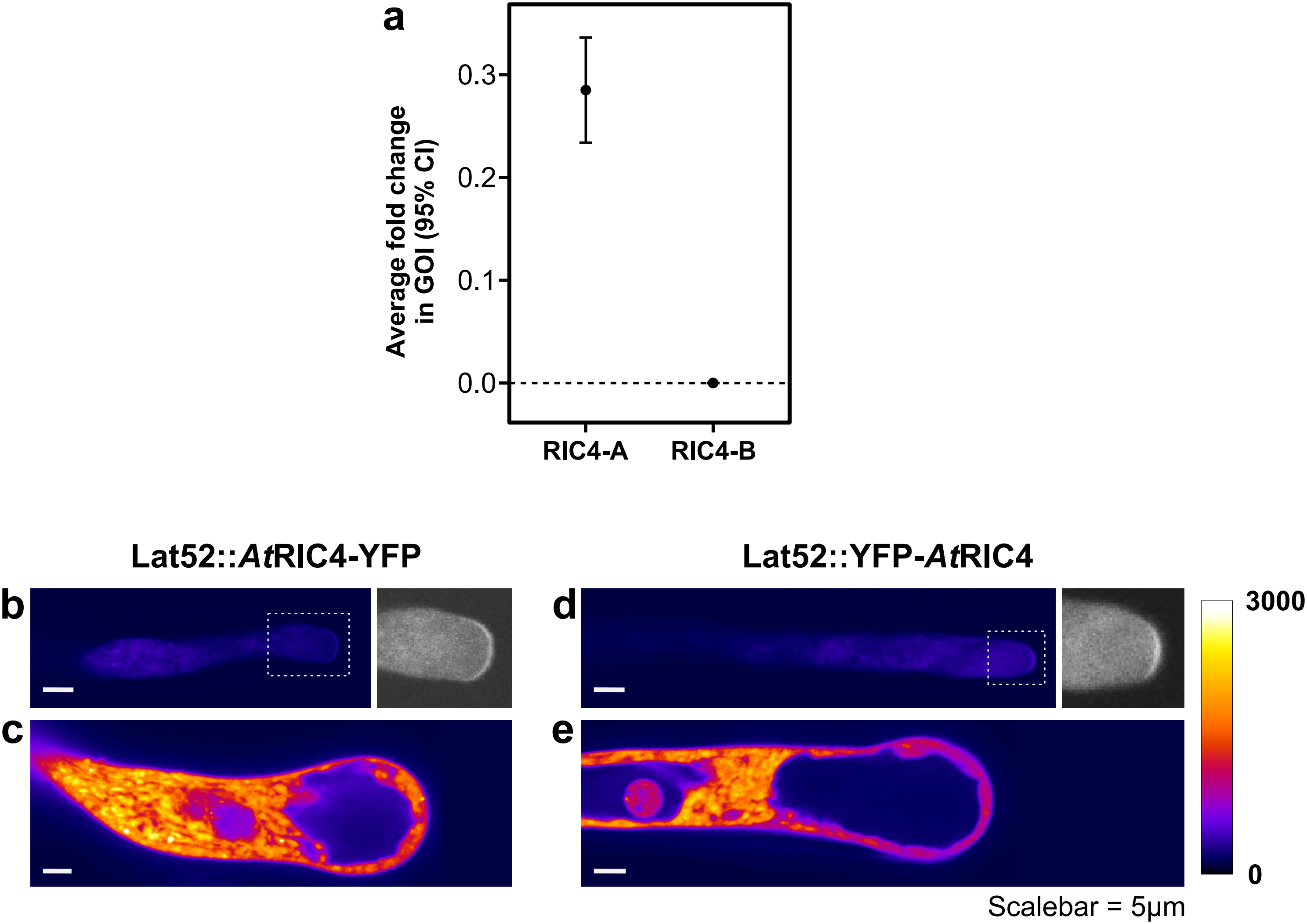

### Supplementary figure 2

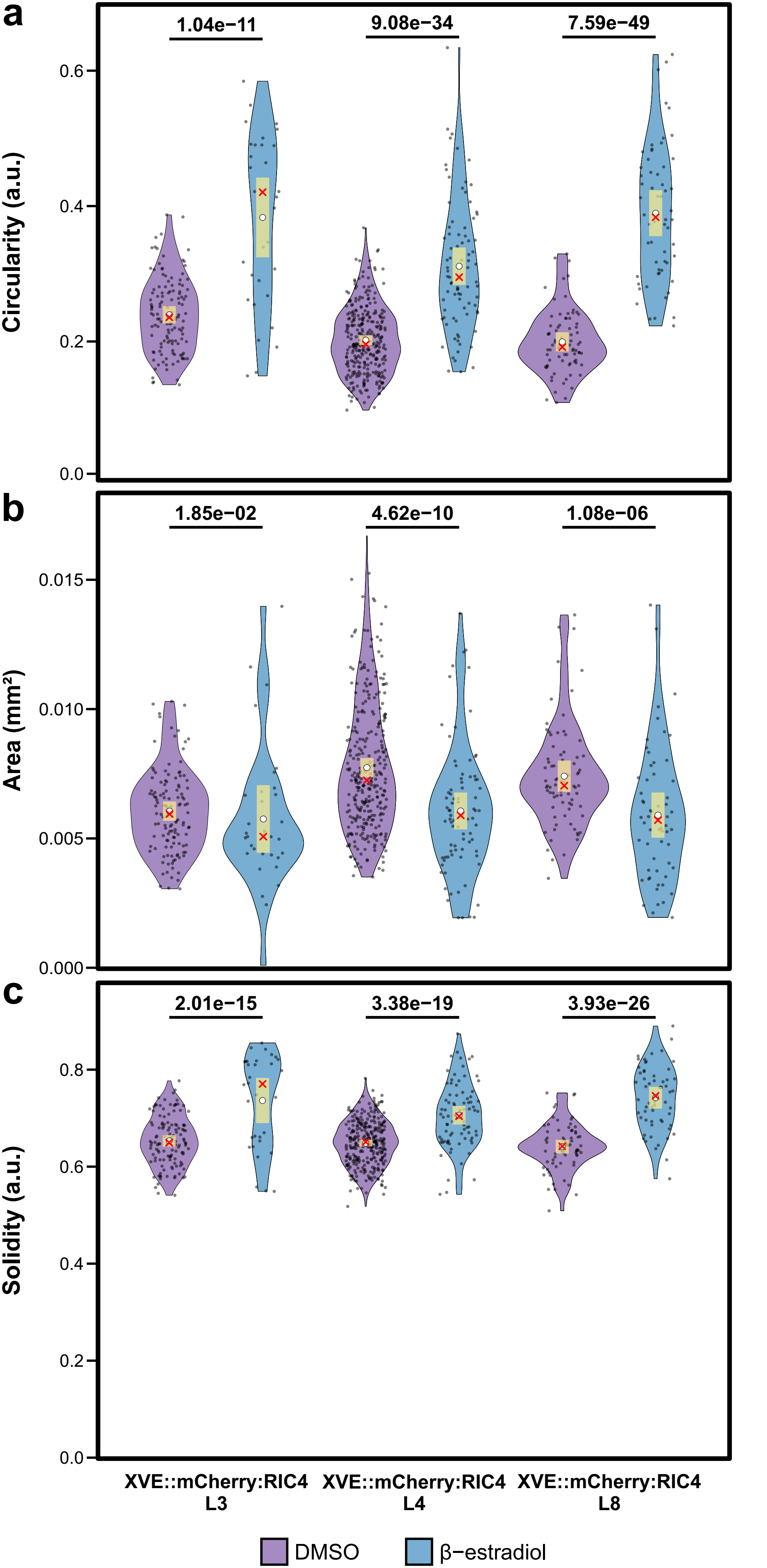

### Supplementary figure 3

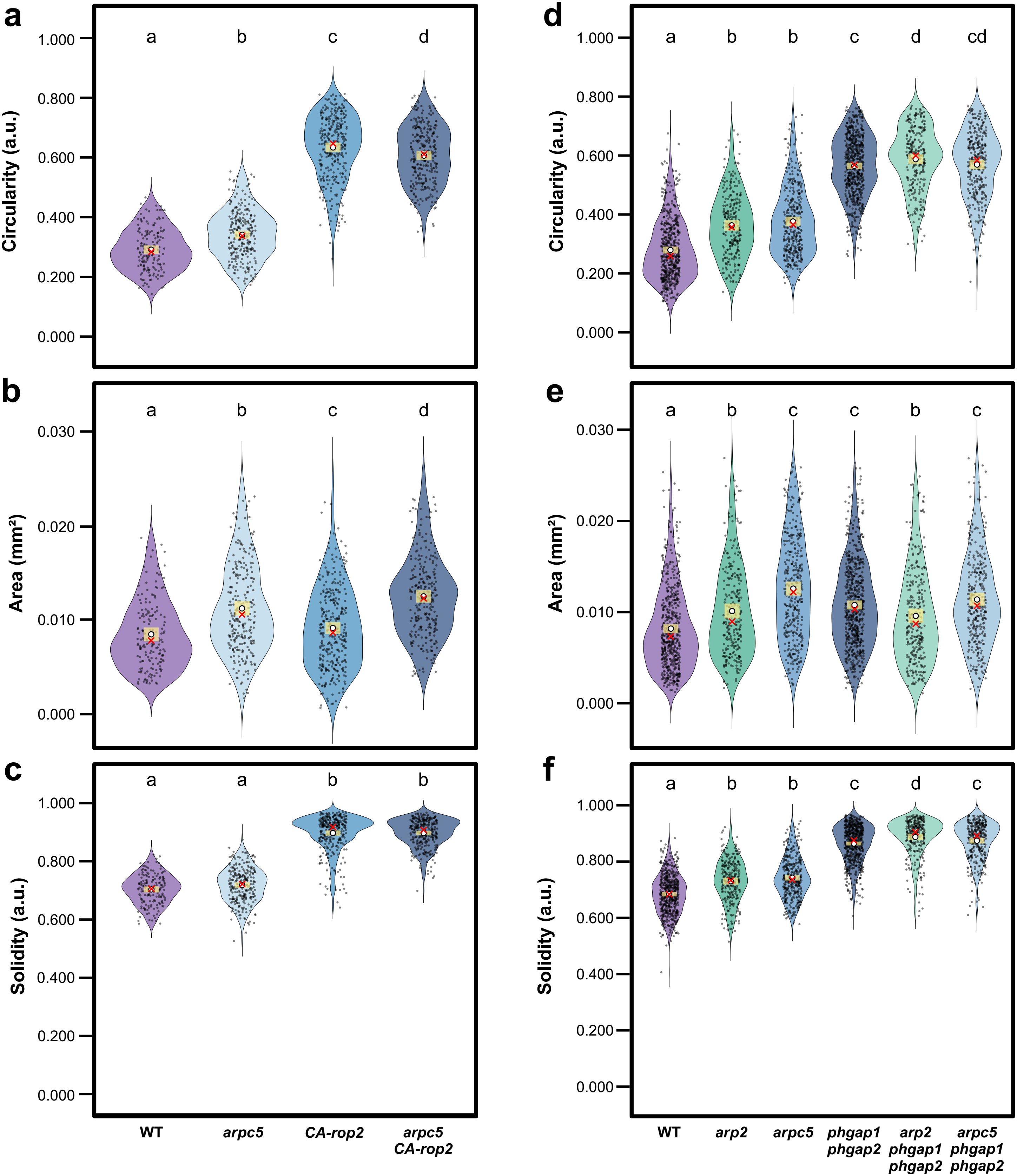

### Supplementary figure 4

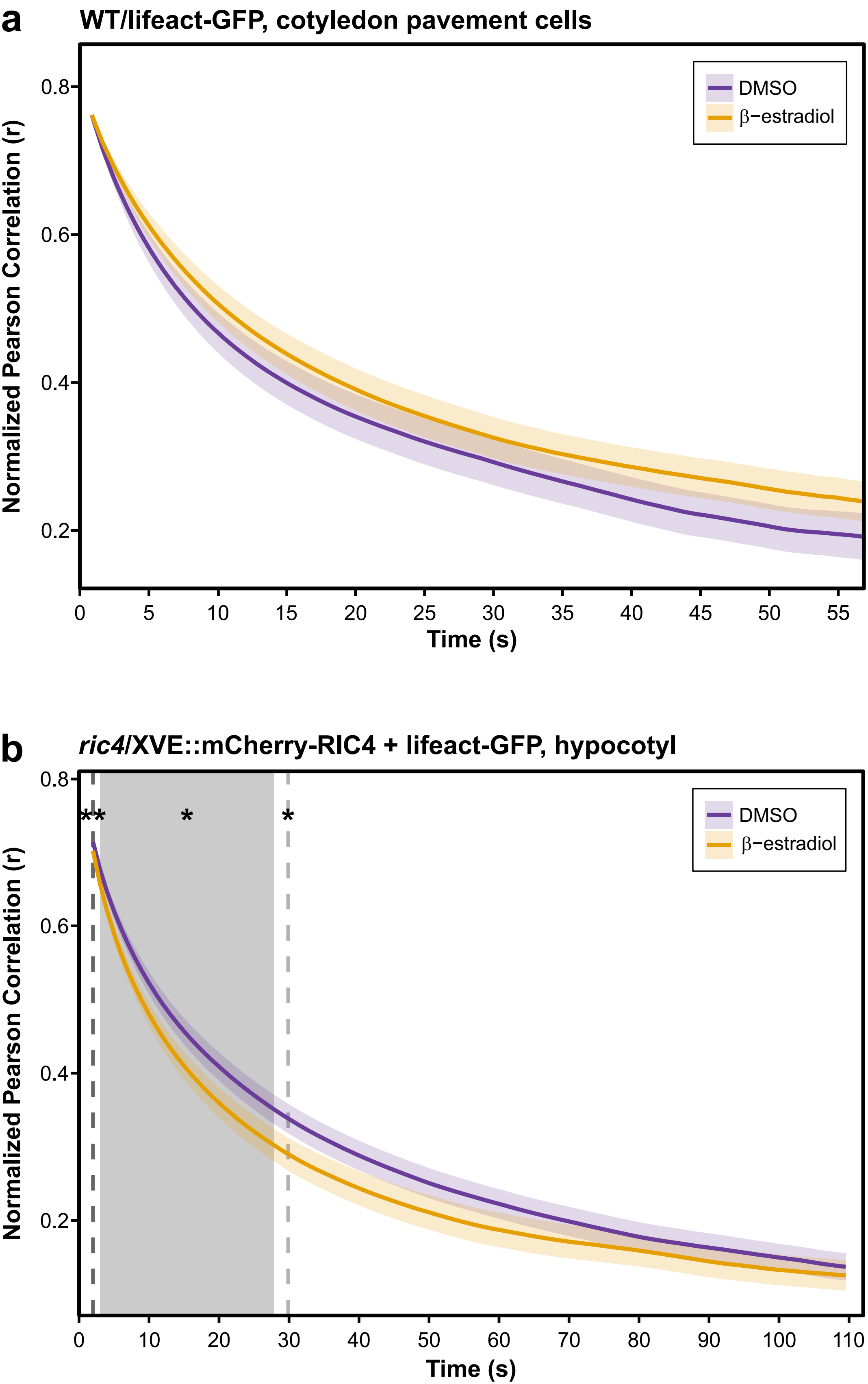
